# Chromatin potential identified by shared single cell profiling of RNA and chromatin

**DOI:** 10.1101/2020.06.17.156943

**Authors:** Sai Ma, Bing Zhang, Lindsay LaFave, Zachary Chiang, Yan Hu, Jiarui Ding, Alison Brack, Vinay K. Kartha, Travis Law, Caleb Lareau, Ya-Chieh Hsu, Aviv Regev, Jason D. Buenrostro

## Abstract

Cell differentiation and function are regulated across multiple layers of gene regulation, including the modulation of gene expression by changes in chromatin accessibility. However, differentiation is an asynchronous process precluding a temporal understanding of the regulatory events leading to cell fate commitment. Here, we developed SHARE-seq, a highly scalable approach for measurement of chromatin accessibility and gene expression within the same single cell. Using 34,774 joint profiles from mouse skin, we develop a computational strategy to identify *cis-*regulatory interactions and define Domains of Regulatory Chromatin (DORCs), which significantly overlap with super-enhancers. We show that during lineage commitment, chromatin accessibility at DORCs precedes gene expression, suggesting changes in chromatin accessibility may prime cells for lineage commitment. We therefore develop a computational strategy (chromatin potential) to quantify chromatin lineage-priming and predict cell fate outcomes. Together, SHARE-seq provides an extensible platform to study regulatory circuitry across diverse cells within tissues.

## Introduction

Regulation of chromatin structure and gene expression underlies key developmental transitions in cell lineages (Novershtern et al., 2011; Shema et al., 2018; Spitz and Furlong, 2012). In recent years, genome-wide profiling of gene expression and chromatin has helped uncover mechanisms of chromatin change at key points of multi-lineage cell fate decisions (Shema et al., 2018; Spitz and Furlong, 2012). Prior studies comparing profiles of purified populations at distinct differentiation states have observed that changes in histone modifications and binding of lineage associated transcription factors (TFs) may precede and foreshadow changes in gene expression creating poised or primed chromatin states that bias genes for activation or repression to alter lineage outcomes (Bernstein et al., 2006; Lara-Astiaso et al., 2014; Rada-Iglesias et al., 2011).

Primed or poised chromatin states are classically defined by the acquisition of specific histone modifications. As one example, deposition of the histone modification H3K4me1 has been shown to prime regulatory elements biasing cells (lineage-priming) for differentiation (Lara-Astiaso et al., 2014; Rada-Iglesias et al., 2011) or immune cell activation (Heinz et al., 2010; Ostuni et al., 2013). However, approaches to analyze primed chromatin states rely on bulk measurements of histone modifications largely restricting analysis to well-defined chromatin states and synchronous cell culture models or stem cell systems with well-defined markers for FACS isolation. Prior work has shown that primed (H3K4me1) and active (H3K4me3) chromatin states reflect a stepwise increase of chromatin accessibility at regulatory elements (Lara-Astiaso et al., 2014). We therefore reasoned that an experimental approach to measure chromatin accessibility and gene expression within the same single-cell may enable identification of primed versus active accessible chromatin, providing a means to identify new mechanisms for chromatin mediated lineage-priming, in new cellular contexts, at single-cell resolution.

We and others have reasoned that methods for combining measurements of different layers of gene regulation within single cells may serve to determine regulators of cell differentiation in tissues and function as sensitive markers of cell identity and cell potential (Kelsey et al., 2017; Shema et al., 2018). Computational methods (Stuart et al., 2019) have had some success in integrating single cell epigenome, transcriptome and protein measurements (Rusk, 2019) profiled separately; however, because these methods assume these distinct measurements align and reflect a common cell identity, they may not be able to correctly recover changes unique to one data type such as chromatin accessibility mediated lineage-priming or lineage-foreshadowing. Emerging single cell “multi-omic” technologies offer a direct means to determine the coordination between layers of gene regulation, including the epigenome and gene expression. Prior studies have sought to correlate gene expression with regulatory element accessibility (Cao et al., 2018; Chen et al., 2019; Zhu et al., 2019). However, these approaches have either limited throughput or limited sensitivity (Cao et al., 2018; Chen et al., 2019; Zhu et al., 2019), prohibiting their ability to recover fine but important distinctions between chromatin accessibility and gene expression.

Here, we investigate the dynamics of the epigenomic and transcriptomic basis of cellular differentiation, by developing Simultaneous High-throughput ATAC (Buenrostro et al., 2013) and RNA Expression with sequencing (SHARE-seq), for individual or joint measures of single-cell chromatin accessibility and gene expression at low-cost and massive scale. Using SHARE-seq, we profiled 84,426 cells across 4 different cell lines and 3 tissue types, including mouse lung, brain, and skin. In particular, applying SHARE-seq to mouse skin shows that cell type definitions are correlated between chromatin accessibility and gene expression, with notable exceptions including high expression variability for cell cycle genes with little to no associated changes in chromatin accessibility. We leverage the heterogeneity across cells to infer chromatin-expression relationships and identify 63,110 peak accessibility-gene expression associations in adult mouse skin. High-density peak-gene associated regions, which we refer to as Domains Of Regulatory Chromatin (DORCs), are enriched for lineage-determining genes and overlap with known super-enhancers (Adam et al., 2015). Strikingly, during hair follicle differentiation, chromatin at DORC-regulated genes become accessible before induction of the corresponding gene’s expression, identifying a role for chromatin accessibility in priming chromatin states. Finally, building upon this finding, we use lineage-priming of chromatin accessibility to predict cellular trajectories during cell differentiation. We develop an analytical framework to systematically compute differences in chromatin and gene expression to predict cell fate choices *de novo* (chromatin potential). Thus, we describe an experimental and analytical basis for integrated measurements of the epigenome and transcriptome enabling new avenues to uncover principles of gene regulation and cell fate specification across single cells in diverse systems.

## Results

### SHARE-seq for joint profiling of chromatin accessibility and gene expression at scale

To create a chromatin accessibility and mRNA expression co-profiling approach that is both scalable and sensitive, we built upon SPLiT-seq (Rosenberg et al., 2018), a combinatorial indexing method for scRNA-seq, to develop SHARE-seq, which uses multiple rounds of hybridization-blocking to uniquely and simultaneously label mRNA and chromatin fragments in the same single cell (**Fig. 1A, Fig. S1A,B, STAR Methods**). Briefly, in SHARE-seq (i) fixed and permeabilized cells or nuclei are transposed by Tn5 transposase to mark regions of open chromatin; (ii) mRNA is reverse transcribed using a poly(T) primer containing a unique molecular identifier (UMI) and a biotin tag; (iii) permeabilized cells or nuclei are distributed in a 96-well plate to hybridize well-specific barcoded oligonucleotides to transposed chromatin fragments and poly(T) cDNA; (iv) hybridization is repeated three times, expanding the barcoding space to approximately 10^6^ (96^3^) barcode combinations (**Fig. S1B, Table S1**), and, following hybridization, cell barcodes are simultaneously ligated to cDNA and chromatin fragments; (v) reverse crosslinking is performed to release barcoded molecules; (vi) cDNA is specifically separated from chromatin using streptavidin beads and each library is prepared for sequencing; and finally, (vii) paired profiles are identified using the common combination of well-specific barcodes (**Fig. S1A**). This barcoding strategy may be extended to even larger experiments, by using additional rounds of hybridization (**Fig. S1B**).

**Figure 1.**
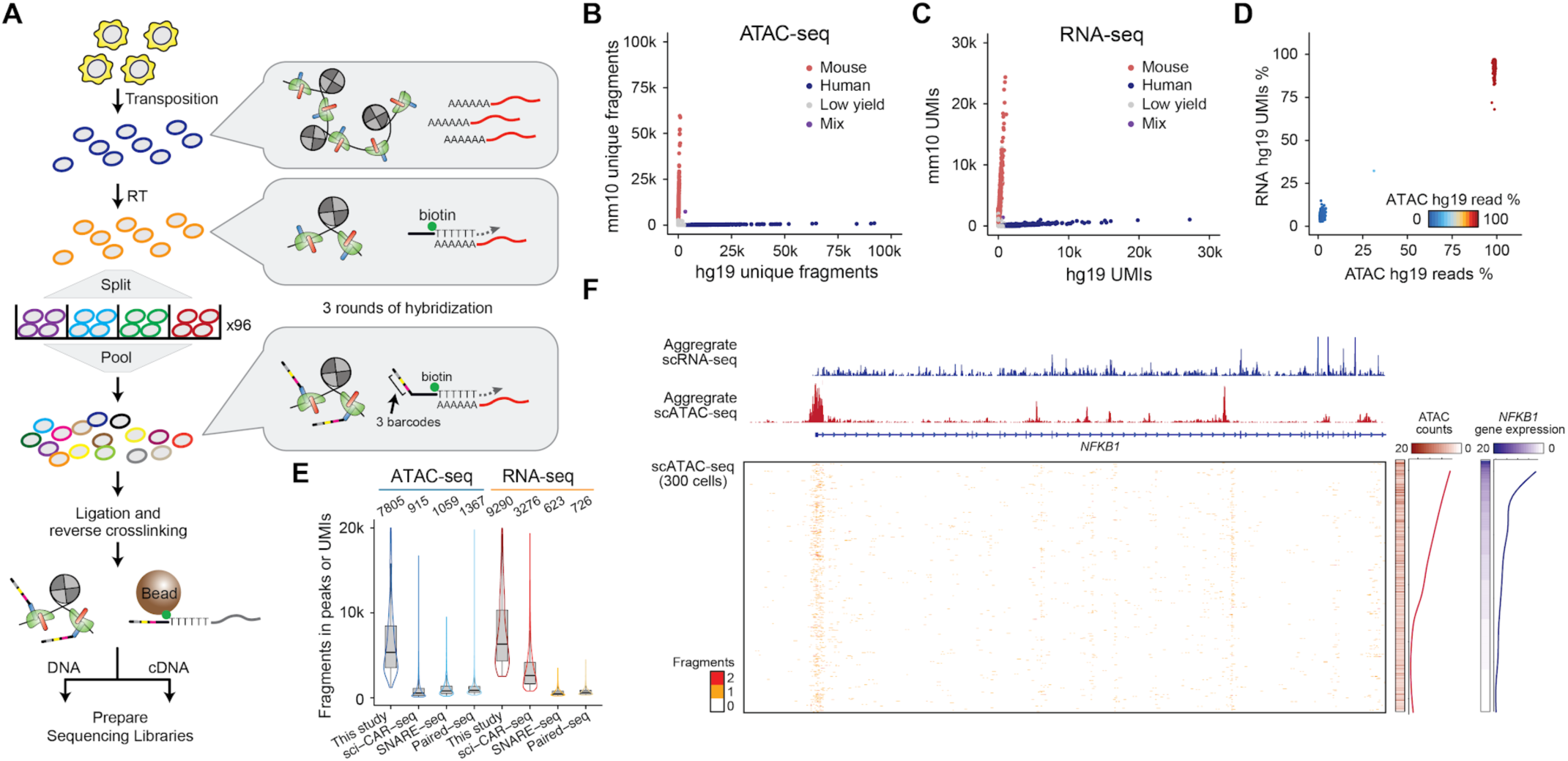
SHARE-seq provides an accurate co-measure of chromatin accessibility and gene expression. **(A)** Workflow for measuring scATAC and scRNA from the same cell using SHARE-seq. **(B,C)** Unique ATAC fragments **(B)**, or RNA UMIs **(C)**, aligning to either the human or mouse genome. The experiment is performed using a mix of human (GM12878) and mouse (NIH/3T3) cell lines. **(D)** The percent of ATAC or RNA reads aligning to the human genome relative to all reads mapping uniquely to the human or mouse genomes. **(E)** Number of ATAC fragments in peaks or RNA UMIs for SHARE-seq (this study), sci-CAR (Cao et al., 2018), SNARE-seq (Chen et al., 2019), or Paired-seq (Zhu et al., 2019). Boxplots denote the medians and the quartile ranges (25 and 75%), length of whiskers represents 1.5 × interquartile ranges (IQRs). **(F)** Aggregated single-cell chromatin accessibility and gene expression profiles in GM12878 cells (top), individual cell profiles (bottom) and single-cell average (right). Single-cells are sorted by the normalized ATAC-seq yield of the depicted *NFkB1* locus.

### SHARE-seq generates high-quality chromatin and expression profiles across diverse cell lines and tissues

To validate specificity and data quality, we first performed SHARE-seq on a mixture of human (GM12878) and mouse (NIH/3T3) cell lines. Human and mouse reads were well separated on both chromatin and transcriptome profiles resulting in 903 human and 1,341 mouse cells passing filter out of 2,000 expected cells (**Fig. 1B-D**). We identified only one cell doublet representing a remarkably low 0.04% collision rate (consistent with the expected rate of 0.052%, **Fig. S1C**), a benefit from the large SHARE-seq barcoding space. Cells passing filter (**STAR Methods**) had on average 2,545 RNA UMIs (9,660 estimated UMI library size) and 8,252 unique ATAC-seq fragments (19,723 estimated library size with 65.5% fragments in peaks) (**Fig. S1D,S1E**).

SHARE-seq had similar performance across replicates and additional cell lines (**Fig. S1F-M**) and showed high concordance with previously published ATAC-seq datasets (**STAR Methods, Fig. S1J**). SHARE-seq also consistently outperformed previously published joint ATAC-RNA approaches (Cao et al., 2018; Chen et al., 2019; Zhu et al., 2019) (**Fig. 1E**), including a technically similar approach (Zhu et al., 2019). Notably, SHARE-seq RNA reads (starting with cells or nuclei) are enriched for intronic regions, similar to single nucleus RNA-seq (snRNA-seq) (Habib et al., 2016) (**Fig. S1N**), which may be due to cell membrane lysis and serial washes used in the protocol. Intronic RNA is enriched for nascent RNA, which can be used not only to identify cell types (Habib et al., 2017), but also to investigate temporal processes within single cells (La Manno et al., 2018). Finally, chromatin accessibility at the *NFkB1* locus and *NFkB1* gene expression significantly co-varied across single-cells (Spearman ρ = 0.31, *p* < 10^−6^, Z-test), validating our expectation that increased chromatin accessibility results in higher gene expression and that SHARE-seq may be used to query chromatin-gene expression relationships (**Fig. 1F**).

Further validating SHARE-seq’s utility, we found that it performed well with cells or nuclei from a broad range of tissues, including mouse skin, brain and lung tissues (**Fig. 2A-C, Fig. S2**). SHARE-seq performed comparably to scATAC-only approaches (Lareau et al., 2019; Mezger et al., 2018) applied to adult mouse lung (**STAR Methods, Fig. 2B**), and to snRNA-seq (Habib et al., 2017) (https://support.10xgenomics.com/single-cell-gene-expression/datasets/2.1.0/nuclei_2k) and scRNA-seq (Saunders et al., 2018; Zeisel et al., 2018) from adult mouse brain (**Fig. 2C** and **Fig. S2D-I**). Importantly, SHARE-seq also enabled experiments at a substantially lower cost than prior methods (**Supplementary Note**). Altogether, these validate the accuracy and utility of this approach for integrated measures of chromatin accessibility and gene expression within cell lines or primary tissues.

**Figure 2.**
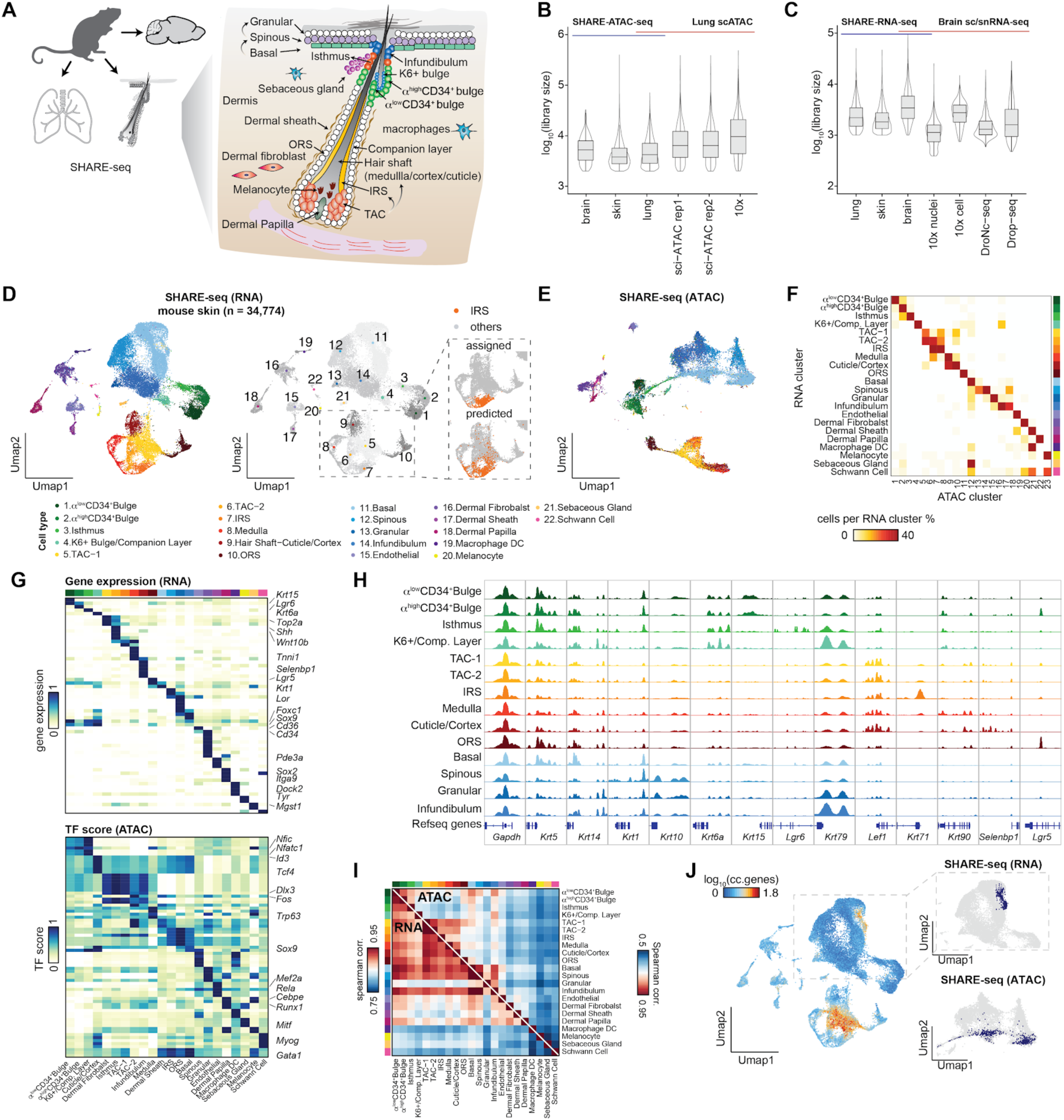
SHARE-seq enables joint profiling of chromatin accessibility and gene expression in tissues. **(A)** A schematic of tissues profiled with SHARE-seq, highlighting the cellular diversity within mouse skin. **(B,C)** Comparison of library size estimates of SHARE-seq and other single-cell or nuclei-based approaches for scATAC-seq **(B)** and scRNA-seq **(C)** approaches. Boxplots denote the medians and the quartile ranges (25 and 75%), length of whiskers represents 1.5× interquartile ranges (IQRs). **(D)** SHARE-seq UMAP visualization of single-cells derived from mouse skin showing UMAP coordinates defined by RNA. Points colored by clusters are defined by RNA clustering, cell types are assigned to clusters on the basis of marker genes, TF motifs, and chromatin accessibility peaks. Computational pairing (Stuart et al., 2019) of scATAC-seq to scRNA-seq (right), colored by predicted cell type. The IRS cluster is highlighted. **(E)** SHARE-seq UMAP visualization of single-cells derived from mouse skin showing UMAP coordinates defined by ATAC. **(F)** Heatmap showing the proportion of cells in the RNA cluster that overlaps with chromatin defined clusters. **(G)** Marker gene expression and TF motif scores for each cluster. **(H)** Aggregated scATAC-seq tracks denoting marker chromatin accessibility peaks for each cluster. **(I)** The cluster-cluster correlation (spearman) of scATAC-seq (top right) and scRNA-seq (bottom left). **(J)** Cells colored by the activity of cell cycle genes (left panel). An RNA cluster marked by high expression of cell cycle genes is highlighted in scRNA UMAP (top right panel) and scATAC UMAP space (bottom right panel).

### Broad congruence between chromatin and RNA defined cell types from SHARE-seq

To utilize SHARE-seq to query the relationship between chromatin accessibility and gene expression, we focused on murine skin. Mammalian skin is enriched for cell types from diverse lineages (including multiple populations of epithelial cells, fibroblasts, immune cells, and neural crest-derived cells) — some are highly proliferative while others are dormant or slow-cycling — with multiple populations of stem cells giving rise to well-defined cell types. Moreover, previous analyses of cellular diversity and chromatin state provide a rich resource to further validate SHARE-seq (Adam et al., 2015; Cohen et al., 2018; Fan et al., 2018; Joost et al., 2018; Lien et al., 2011; Salzer et al., 2018). Thus, we reasoned the skin may be an ideal tissue to resolve cellular and regulatory diversity across cells at different proliferation or differentiation status (Hsu et al., 2014).

Leveraging the increased throughput and resolution of SHARE-seq, we assessed the congruence between the epigenome and transcriptome across an atlas of 34,774 high-quality profiles from adult mouse skin (**Fig. 2D, Fig. S3A,B**). To define cell subsets, we clustered the RNA portion of SHARE-seq data and identified consensus expression signatures for each cluster, highlighting known and novel markers (**Table S2**). We projected the cells based on the ATAC-seq and RNA-seq independently to a low dimensional space using UMAP (**STAR Methods**), and found that both projections separated these scRNA-seq defined clusters (**Fig. 2D,E**). SHARE-seq not only resolved cell types from distinct lineages (epithelium, fibroblasts, melanocytes, immune cells), but could also distinguish between cells of closely related types (for example, α^high^ CD34^+^ bulge vs. α^low^ CD34^+^ bulge (Blanpain et al., 2004)). Moreover, cell membership in subsets identified by scATAC-seq was highly congruent with their membership within scRNA-seq clusters (**Fig. 2F, Fig. S3C**), and both measures revealed the same major cell type classes, such as transit-amplifying cells (TACs), inner root sheath (IRS), outer root sheath (ORS), and hair shaft cells (**Fig. 2D-F**).

The cells within the RNA-based clusters can also be distinguished by chromatin accessibility features, further confirming their identity (**Fig. 2G,H**). We annotated clusters by the activity of lineage-determining TFs, which we inferred by TF activity scores from the scATAC-seq data (**Fig. 2G**) (Schep et al., 2017), and their correlation to TF expression levels (**Fig. S3D-F, STAR Methods**). This analysis revealed global transcriptional activators *Dlx3* (Adam et al., 2015) (a hair follicle differentiation gene) and *Sox9* (Adam et al., 2015) (a key regulator of the hair follicle stem cell identity), and repressors *Zeb1* (Spaderna et al., 2008) and *Sox5* (Huang et al., 2008), among many others (**Fig. S3F,G**). Thus, we found SHARE-seq provides insight into cell identity at multiple scales, including chromatin regulation by key lineage-determining TFs, enabling the analyses of cellular and regulatory atlases at scale.

Nevertheless, some cell subsets (*e.g.*, in differentiation) or states (*e.g.*, cell cycle) may be identified at higher resolution by either chromatin or gene expression features. Specifically, grouping clusters by their aggregate (pseudo-bulk) profiles more distinctively revealed chromatin accessibility differences between permanent portion (clusters 1-4) and regenerative portion (clusters 5-9) of hair follicle. Conversely, cells corresponding to the granular layer are easier to distinguish as a unique cluster at the gene expression level (**Fig. 2I** and **Fig. S3C**). Moreover, a subset of actively proliferating basal cells strongly expressing cell-cycle genes (**Fig. 2J**) was not identified as a coherent cluster by chromatin accessibility (with either of four different dimensionality reduction approaches, **Fig. 2J, Fig. S3H-J, STAR Methods**).

We reasoned that SHARE-seq can be used to directly test the accuracy of computational approaches (Stuart et al., 2019) that pair data from scATAC-seq and scRNA-seq from separately measured cells; such methods typically assume congruence, and may miss asynchrony or distinctions between these features of cellular identity. We thus tested a Canonical Correlation Analysis (CCA)-based method (Stuart et al., 2019) by providing the computational approach the scATAC-seq and scRNA-seq portions of the SHARE-seq measurements separately, and comparing its inferred pairing (defined as membership in the same cluster) to the correct (measured) coupling. Profiles from the same cell were properly assigned to the same cell subset with variable accuracy (74.9% in skin and 36.7% in mouse brain) (**Fig. S4**), with most mis-assignments between clusters representing similar cell types (*e.g.*, IRS to TACs, **Fig. 2D**). The mis-assignments may result from chromatin changes preceding or succeeding gene expression during differentiation from TACs to IRS, as we discuss below. Such computational errors may be exasperated in differentiating cell types as seen in the skin or across highly diverse populations as seen in the brain. Together, this suggests SHARE-seq may help either train computational pairing approaches across tissues or test their performance and help with further improvements.

### Paired measurements associate chromatin peaks and target genes in *cis*

Cells exhibit significant variation in both gene expression (Marinov et al., 2014) and the underlying regulation of chromatin (Buenrostro et al., 2015), due to both intrinsic (*e.g.*, bursts of expression (Larsson et al., 2019)) and extrinsic (*e.g.*, cell size, level of regulatory proteins (Lin and Amir, 2018)) factors. We reasoned that joint measurements in SHARE-seq may allow us to infer the relationship between variation in chromatin and gene expression. To test this, we developed an analytical framework to link distal peaks to genes in *cis*, based on the co-variation in chromatin accessibility and gene expression levels across cells, while controlling for technical biases in chromatin accessibility measurements (**Fig. 3A, STAR Methods**). We first applied this approach to a data set of 23,278 GM12878 cells, and identified 13,277 significant peak-gene associations (**Fig. S5**, *p* < 0.05, FDR = 0.11). Importantly, we found down-sampling of either cell numbers or number of detected reads (matching library quality to those of previous chromatin/RNA reports (Cao et al., 2018)) dramatically reduces the ability to discover peak-gene associations (**Fig. 3B, Table S3**).

**Figure 3.**
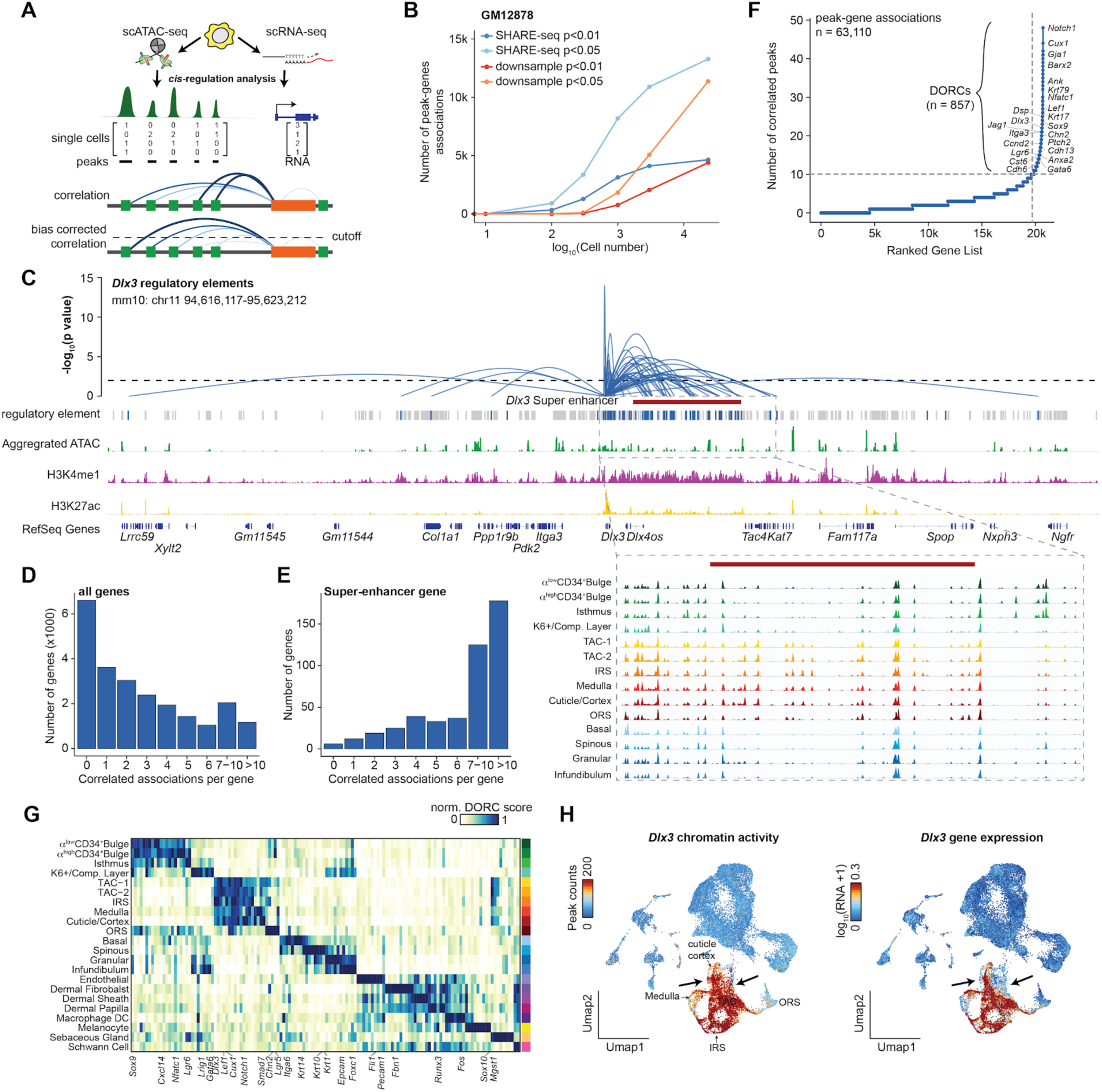
*Cis-*regulation determines Domains Of Regulatory Chromatin (DORCs). **(A)** Schematic depicting an analytical framework for analysis of distal regulatory elements and expression of genes. **(B)** Number of peak-gene associations after down-sampling the number of cells or reads within the GM12878 SHARE-seq data set. Reads are down-sampled to match the number of reads recovered to match those obtained by sci-CAR (Cao et al., 2018). **(C)** Loops denote the *p*-value of chromatin accessibility of each peak and *Dlx3* RNA expression. Loop height represents the significance of the correlation. H3K4me1 and H3K27ac ChIP-seq tracks and super-enhancer annotation generated from isolated TAC population (Adam et al., 2015). **(D, E)** The number of significant peak-gene associations for all genes **(D)** and previously defined (Adam et al., 2015) super-enhancer genes **(E). (F)** The number of significantly correlated peaks (*p* < 0.05) for each gene. Known super-enhancer regulated genes are highlighted. **(G)** Representative DORCs for each defined cluster, values are normalized by the min and max activity. **(H)** The peak counts of all *Dlx3* correlated peaks (left) and *Dlx3* gene expression (right) colored in UMAP. The arrows point to regions with differential signals.

Applying this framework to murine skin dataset, we identified 63,110 significant peak-gene associations (within ±50kb around transcription start sites (TSSs), *p* < 0.05, FDR = 0.1, after filtering peaks associated with multiple genes, **Table S4**). These peak-gene associations were enriched proximally to the TSS (**Fig. S6A,B**, *p* < 2.2×10^−16^, KS-test). We found, only 10,154 additional peaks were associated with more than a single gene (**Fig. S6C,D**) suggesting most regulatory elements (83.9%) only regulate a single gene. Interestingly, most of the chromatin peaks (82%, **Fig. S6E**) were not correlated with the expression of any gene, a finding that may support a previous observation that only a small portion of candidate enhancers significantly alter the expression of genes (Gasperini et al., 2019).

A subset of genes, including key fate-determination genes, were associated with a large number of peaks (*p* < 2.2×10^−16^, permutation test, **STAR Methods**). For example, 83 and 53 peaks were significantly associated (within ±500kb around TSSs, *p* < 0.05) with *Dlx3*, highly expressed in TACs (**Fig. 3C**), and *Cxcl14*, highly expressed in bulge (**Fig. S6F**), respectively (Adam et al., 2015). These results are reminiscent of previous observations describing regulatory locus complexity at key lineage genes (González et al., 2015). Further, we found regions with high-density of peaks-gene associations significantly overlap known super-enhancers (Adam et al., 2015) (**Fig. 3C, Table S5**, 2.1 fold enrichment, *p* = 10^−238^, hypergeometric test) — enhancer regions that are cell-type specific and highly enriched in histone H3K27 acetylation (Whyte et al., 2013). This relationship was not simply driven by super-enhancer length (Spearman ρ = 0.04; **Fig. S6G**). Furthermore, super-enhancers regulated genes are associated with more peaks compared to all genes (10.9 *vs*. 4.4 associated peaks per gene on average *p* < 2.2×10^−16^, KS-test, **Fig. 3D,E, Fig. S6H**). Notably, while peak-gene associations are enriched at known super-enhancers, we find that densely regulated genes also exhibit interactions outside the annotated super-enhancer which may reflect the false discovery of our approach (FDR = 0.1) or limitations in calling super-enhancers using ChIP-seq which often does not incorporate the 3D configuration of the locus (Schoenfelder and Fraser, 2019). Finally, most annotated cell cycle genes (n = 97) had lower than expected number of peak-gene interactions (on average 3.4 interactions for cell cycle genes *vs*. 4.4 interactions for all genes; *p* = 0.026, *t*-test), further supporting a limited contribution of chromatin accessibility to cell cycle associated gene expression and suggesting that variable expression is not sufficient for determining peak-gene associations.

### Domains Of Regulatory Chromatin (DORCs) identify key lineage-determining genes *de novo*

We define the 857 regions with an exceptionally large (>10) number of significant peak-gene associations as “Domains Of Regulatory Chromatin” (DORC), identified as those exceeding an inflection point (“elbow”) when ranking genes by the number of significant associations (**Fig. 3F**). Genes associated with super-enhancers were strongly enriched in DORC-regulated genes (*p* = 10^−97^, hypergeometric test). We quantified the activity of DORCs as the sum of accessibility at peaks significantly associated with the DORC-regulated gene. DORC accessibility scores were highly cell-type specific, and DORC-regulated genes were identified when conducting the analysis within a cell type (GM12878 cells) or across diverse cells types in tissues (**Fig. S5**). The DORCs identified within sub-populations strongly overlap with DORCs identified with all cells (*p* = 10^−201^, hypergeometric test, **Fig. S6I**). Moreover, chromatin accessibility of DORCs were strongly enriched for known key regulators of lineage commitment across the expected lineages (**Fig. 3G, Fig. S6J**). For example, *Sox9*, a master regulator of stem cell fate commitment (Nowak et al., 2008), is a DORC-regulated gene, and this DORC has high activity in stem cell populations (**Fig. 3G**).

There were significant differences between DORCs even in closely related populations, suggesting DORCs may help predict novel regulators that distinguish them. Interestingly, in some cases, high DORC activity in a particular cell subset presented little to no gene expression of the DORC-regulated gene, suggesting a gain of chromatin accessibility does not always equate to productive transcription. For example, while *Dlx3* DORC activity and *Dlx3* gene expression were generally correlated in TAC/IRS/Hair shaft cells, this was not the case within cuticle/cortex cells (**Fig. 3H, Fig. S6K**). Thus, DORCs provide an unsupervised, readily accessible approach to simultaneously identify key lineage-determining genes and the regions that regulate them at single-cell resolution, without the need to know the cell type identity in advance, isolate cell subsets, and conduct challenging ChIP-seq experiments from primary samples (see **Discussion**).

### Lineage priming at enhancers precedes gene expression in DORCs

The hair follicle is a highly regenerative epithelial tissue that cycles between growth (anagen), degeneration (catagen), and rest (telogen). At the anagen onset, hair follicle stem cells located at the bulge and hair germ proliferate transiently to produce the short-lived TACs. These TACs are one of the most proliferative cells in adult mammals — they proliferate rapidly to produce multiple morphologically and molecularly distinct downstream differentiated cell types that constitute the mature hair follicle, including the companion layer, IRS (Henle’s layer, Huxley’s layer, IRS cuticle) and hair shaft (hair shaft cuticle, cortex, medulla) (Zhang and Hsu, 2017; Zhang et al., 2016). Previous studies have shown that TACs display molecular heterogeneity but still maintain a degree of lineage plasticity (Xin et al., 2018; Yang et al., 2017). The unique features of TACs provide an interesting context to study chromatin-expression relationships in cells that are required to dynamically change their epigenome to choose lineage fates, while undergoing rapid proliferation.

We readily recovered hair follicle differentiation trajectories from chromatin accessibility (**Fig. 4A**), whereas similar analysis of the RNA profile led to challenges, due to the strong expression of cell cycle genes in the rapidly proliferating TACs (**Fig. 2I, Fig. S7**). Single cell chromatin profiles were ordered into three lineage trajectories, IRS, medulla, and cuticle/cortex, differentiated from TACs (**Fig. 4A**). The detailed structures in IRS (including Henle’s layer, Huxley’s layer, and the IRS cuticle) were not fully resolved due to the rareness of these cell types (672 (∼2%) IRS cells out of 34,774 cells, consistent with previously reported 1-3.8% cell types (Joost et al., 2016), though we did identify a subset of cells located between the IRS and medulla lineages (**Fig. 4A**), which may suggest distinct differentiation routes to these IRS subsets, as we explore below.

**Figure 4.**
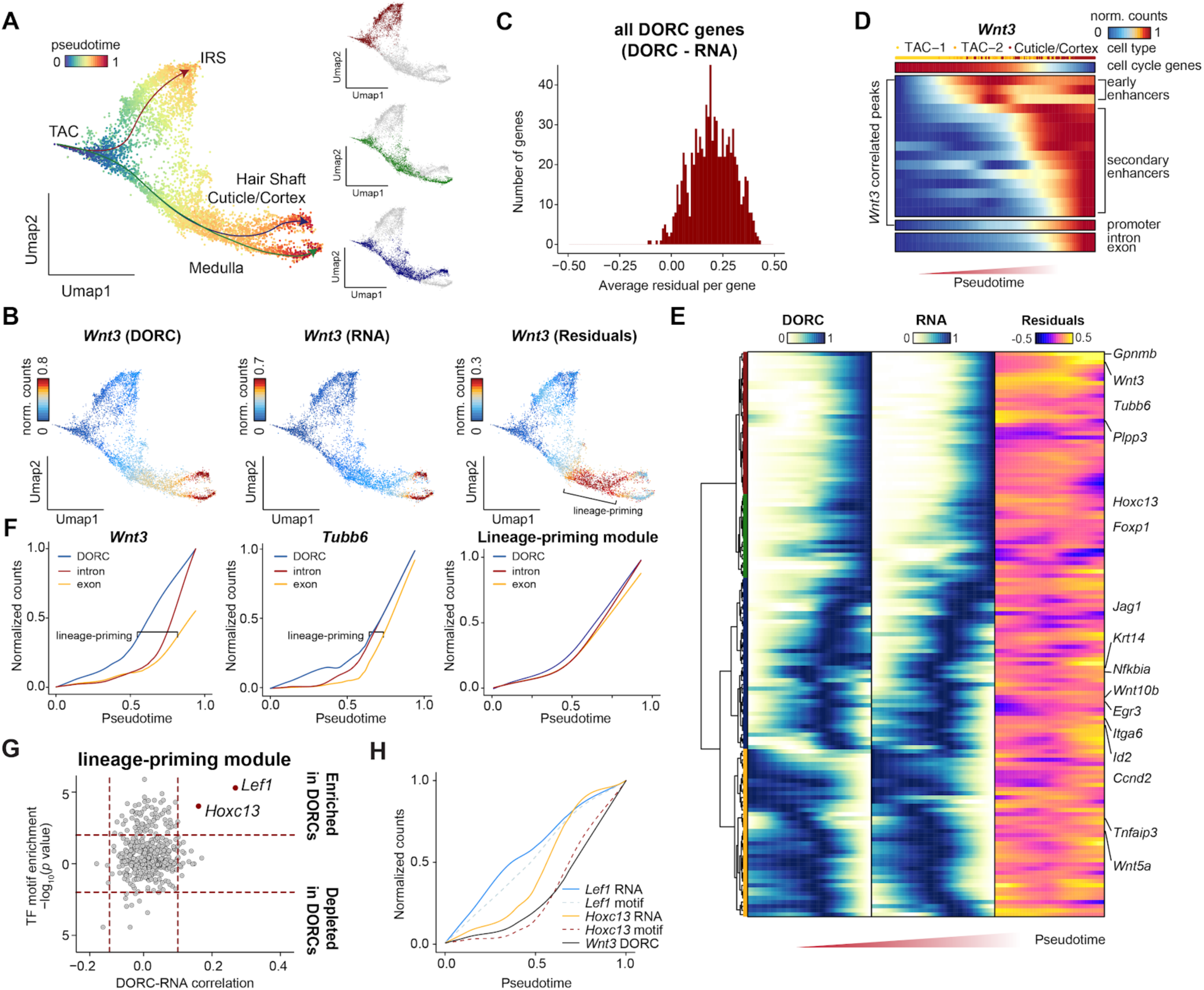
Lineage dynamics of chromatin and expression defines lineage priming. **(A)** Pseudotime for three cell fate decisions shown on scATAC UMAP coordinates. **(B)** Difference (residuals) for *Wnt3* between chromatin accessibility and gene expression for the regenerative portion of the hair follicle. **(C)** Histogram of the average difference (residuals) for each gene between chromatin accessibility and gene expression. **(D)** Dynamics of gene expression (intron and exon) and individual chromatin accessibility peaks for the Cuticle/cortex lineage. **(E)** Hierarchical clustering of chromatin accessibility, expression of DORC-regulated genes and the difference between chromatin accessibility and gene expression (residuals) for the cuticle/cortex lineage. Cells are ordered by pseudotime. **(F)** Lineage dynamics for individual DORC-regulated genes highlighting lineage-priming in *Wnt3* (left), *Tubb6* (middle), and the mean of the lineage-priming module (red cluster in panel **E**). **(G)** TF motif enrichment in lineage-priming DORCs plotted against Spearman correlation of the lineage-priming module DORC score and gene expression of individual TFs. **(H)** Lineage dynamics of *Lef1* and *Hoxc13* motif scores and gene expression precede *Wnt3* DORC activation in the hair shaft lineage.

Systematically analyzing the onset of accessibility and gene expression along differentiation pseudotime from TACs to cuticle/cortex cells revealed that DORCs generally become accessible prior to the onset of their associated gene’s expression, consistent with lineage-priming. Given their particular association with lineage-determining TFs, we hypothesized that DORCs may play an important role in differentiation. For example, *Wnt3* RNA became detectable at the late stage of hair shaft differentiation, consistent with previous findings (Millar et al., 1999). However, accessibility in the *Wnt3* DORC activated at TACs prior to gene expression before lineage commitment (**Fig. 4B**), which we quantified by computing “residuals” (defined as the difference of chromatin accessibility and expression of the gene). Systematic analysis typically found positive residuals across all DORC-regulated genes and lineages (**Fig. 4C, Fig. S8A**), despite the overall correlated accessibility of DORCs and the expression of the DORC-regulated genes (by definition, **Fig. S8B**). Thus, sufficiently high RNA expression may be detectable only within a subset of cells with accessible chromatin at that gene’s locus.

To further understand the possible underlying cause of these residuals, we tracked the changes in accessibility in individual peaks in the *Wnt3* DORC along differentiation pseudotime from TACs to cuticle/cortex cells (**Fig. 4D**). We found a sequential activation of peaks, with individual enhancer peaks activating much earlier (in pseudotime) than the *Wnt3* promoter, followed by activation of nascent RNA expression (estimated by intron counts) and finally mature RNA expression (estimated by exon counts) (**Fig. 4D**). This pattern of peak activation in enhancers prior to expression is apparent across many but not all genes, which we refer to as the “lineage-priming module” defined by sharing similar residuals (**Fig. 4E, Fig. S8C**). To more directly quantify these differences, we calculated a lag of 0.20 or 0.13 pseudotime units between the respective onset of accessibility in the *Wnt3* or *Tubb6* DORCs and the onset of expression of these genes, first at pre-mRNA and then by mRNA (**Fig. 4F**). Notably, some TACs at late pseudotime are still proliferative (estimated by cell cycle gene expression); however, they already show activated enhancer peaks, suggesting the proliferation and differentiation switch transitioning from TACs to cuticle/cortex lineage (**Fig. 4D**). Altogether, these analyses support the long-standing hypothesis that enhancer activation foreshadows expression of target genes (Lara-Astiaso et al., 2014; Rada-Iglesias et al., 2011) and implicates chromatin accessibility as a marker for lineage-priming (Olsson et al., 2016).

We further investigated the mechanisms leading to chromatin accessibility primed chromatin states and hypothesized that TFs that prime are distinct from TFs that activate enhancers, suggesting a subset of specific TFs may induce lineage-priming. Indeed, we found that binding sites for *Lef1* and *Hoxc13* TFs are strongly enriched (*p* < 10^−4^, KS-test, **Fig. 4G**) in hair shaft lineage-priming DORCs (including the *Wnt3* DORC). Gene expression and TF motif activity (inferred from ATAC-seq) of *Lef1*, a known regulator of the *Wnt* signaling pathway (Clevers, 2006), activated prior to *Hoxc13 (Godwin and Capecchi, 1998)*, implicating *Lef1* as the lineage-priming TF (Merrill et al., 2001). This was followed by expression of *Hoxc13 (Godwin and Capecchi, 1998)*, likely inducing *Wnt3* DORC accessibility and promoting *Wnt3* gene expression (**Fig. 4H, Fig. S8D**). Together, this supports a model, whereby distinct modes of regulation exist to prime chromatin accessibility and foreshadow lineage choice.

### Chromatin potential describes chromatin-to-gene expression dynamics during differentiation

Empowered by our findings, we hypothesized that lineage priming by chromatin accessibility may foreshadow gene expression and may be used to predict lineage choice prior to lineage commitment. To explore this possibility, we focused on DORC-regulated genes, which encompass lineage-determining genes and coincide with strong chromatin signals. We devised an approach to calculate “chromatin potential”, defined as the future RNA state most compatible with a cell’s current chromatin state. To calculate chromatin potential, we first address data sparsity, by smoothing each cell by computing a *k*-nearest neighbor graph (*k*-NN defined by chromatin state, *k* = 50) and averaging chromatin and expression profiles for cells in this neighborhood. Next, we computed RNA-chromatin neighbors (*k-*NN, *k*=10) whereby we find, for each cell (cell *x*, chromatin neighborhood), 10 cells (cell *y*, RNA neighborhood) whose RNA expression of DORC-regulated genes is most correlated to the current chromatin state. Chromatin potential (arrow) is the direction and distance between each cell (cell *x*, chromatin neighborhood) and 10 nearest cells (cell *y*, RNA neighborhood) in chromatin low dimensional space (**Fig. 5A, Fig. S8E,F, STAR Methods**). We note here that this analysis did not rely on the inferred pseudotime. Chromatin potential relates a potential “future” RNA state (observed in another cell) which is best predicted by the chromatin state of a given cell.

**Figure 5.**
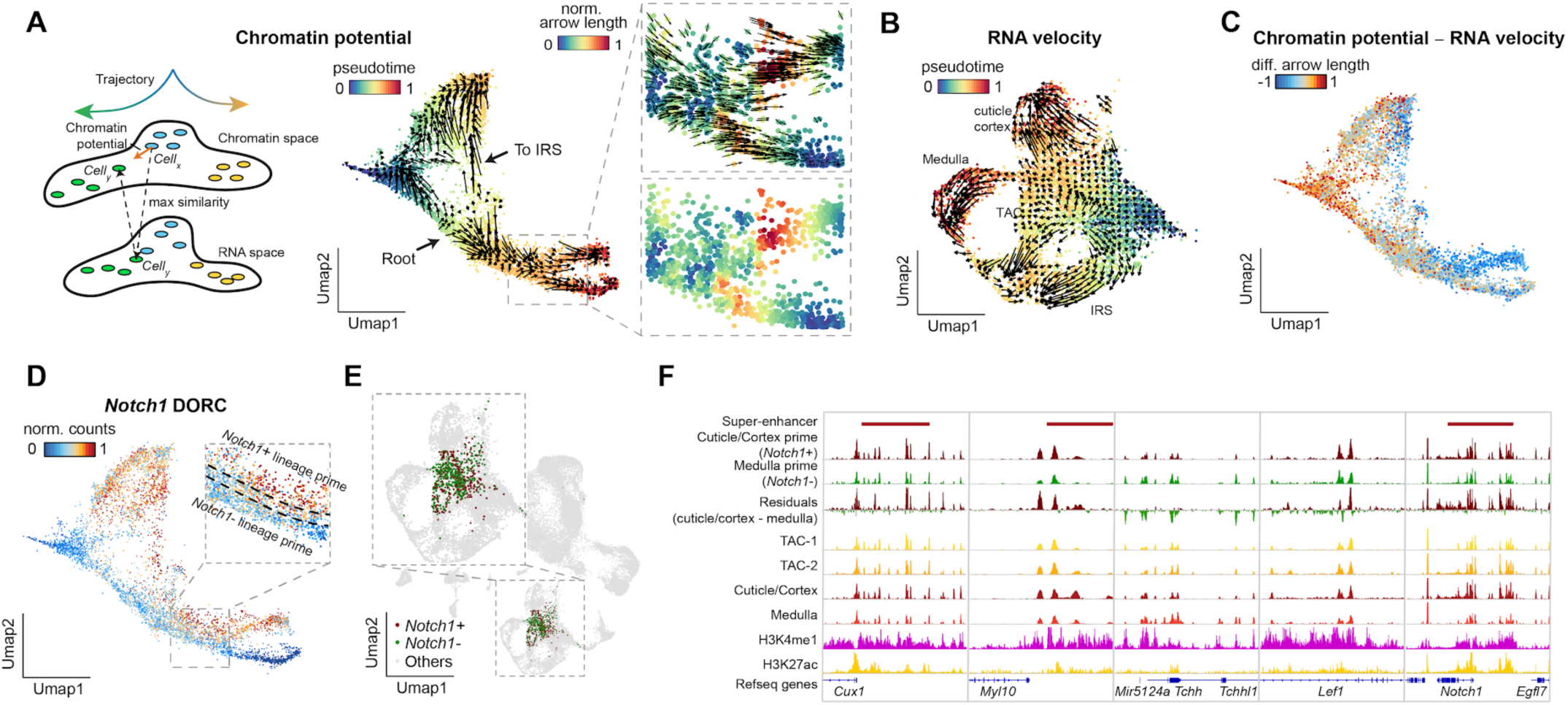
Chromatin potential describes chromatin-to-gene expression dynamics during differentiation. **(A)** Schematic of the conceptual workflow for determining chromatin potential (left). Chromatin potential visualized on the scATAC UMAP space, arrows denote the extrapolated gene expression state of the cell (right). **(B)** RNA velocity visualized on scRNA UMAP coordinates. **(C)** The difference between the neighborhood predicted by chromatin potential and RNA velocity. **(D)** Chromatin accessibility of the *Notch1* DORC, highlighting the lineage-priming region. **(E)** Distribution of *Notch1+* and *Notch1-* lineage primed cells in the scRNA UMAP. **(F)** Aggregated chromatin accessibility profiles of lineage primed cells (*Notch1*+/-), progenitor cells (TACs), and differentiated cells (cuticle/cortex, medulla).

In general, chromatin potential flows from progenitor cells (TACs) to differentiated cells (IRS/Hair shaft). Long arrow length represents a chromatin state reflecting a more differentiated transcriptome. Regions of long arrows suggest lineage commitment at these lineage events occurs as a switch rather than as a gradient (Yang et al., 2017). Chromatin potential is higher at key multi-lineage defining transitions, including the branch point that defines the cuticle/cortex and medulla lineages.

In many key developmental transitions, chromatin potential exceeds our ability to predict future RNA states from the cell’s current RNA state, by either its mRNA or its nascent RNA (as shown by RNA velocity (La Manno et al., 2018)), emphasizing the longer time scales foreshadowed by chromatin states. This is clear by several different measures. First, the “future” RNA state predicted by chromatin potential extends significantly further than that predicted with the current RNA state (**Fig. S8G-I**). Second, RNA velocity derived vectors, which use intronic RNA as a measure of nascent transcription to determine future states, validated our end-point trajectories; however, they provided little resolution of cell fate dynamics within TACs (**Fig. 5B**) (La Manno et al., 2018). In particular, when we predict for each cell its RNA velocity (from RNA) and its chromatin potential (from chromatin-RNA), cells reflecting the “future” RNA state (defined by RNA velocity) results in substantially less chromatin-RNA correlation when compared to the “future” state predicted by chromatin potential (*p* < 2.2×10^−16^, *t*-test, **Fig. S8J**). The discrepancy between RNA velocity and chromatin potential is most prominent in TACs (**Fig. 5C**). Interestingly, chromatin potential has longer reach (prediction timescales) at early stages, whereas RNA velocity (extended to a *k*-NN neighborhood) has further reach (longer arrows) at late pseudotimes (*p* < 2.2×10^−16^, KS-test, **Fig. S8K**). Generally, chromatin potential supported the measure of pseudotime; however, it identified a distinct root-like position, which suggests either an alternative lineage origin or plasticity and route reversal (**Fig. 5A**). Chromatin potential also suggested an alternative route to the IRS which is not identified by RNA velocity (**Fig. 5B**); however, additional experiments using lineage tracing are needed to better elucidate the dynamics of this transition. Thus, chromatin potential allows us to relate the chromatin state of one cell to future RNA states not yet realized in that cell, and to span longer time scales especially in early developmental transitions (see **Discussion**).

Finally, we sought to see how early we could identify markers of lineage commitment, searching for genes whose chromatin state foreshadows lineage commitment far preceding the lineage choice as reflected in their RNA state. To investigate this, we identified DORCs that were differentially active between cuticle/cortex and medulla cells preceding the lineage decision, including *Notch1, Cux1*, and *Lef1* (**Fig. S8L**). Interestingly, *Notch1* is highly expressed in hair shaft cells, whose DORC encompasses 79 peaks. *Notch1* is critical in controlling hair follicle differentiation and acts non-autonomously to regulate the formation of hair shaft and IRS (Pan et al., 2004). The activation of *Notch1* DORC activity coincides with longer arrows associated with cuticle/cortex lineage commitment (**Fig. 5A**). When we partition the lineage-priming region into 3 sub-regions by the DORCs’ accessibility (**Fig. 5D**), *Notch1*^*+*^ and *Notch1*^*-*^ regions showed distinct chromatin patterns with coordinated changes in gene expression, whereas *Notch1*^+^ cells were not distinctly identified by their gene expression pattern alone (**Fig. 5E, Fig. S8L,M**). *Notch1*^+^ and *Notch1*^-^ regions showed chromatin potential to differentiate into cuticle/cortex and medulla lineages, respectively. Consistently, we observed clear chromatin-gene expression differences (residuals) at multiple loci in the lineage-priming region using aggregated genome tracks (**Fig. 5F**). A similar differential trend was observed between fully differentiated cells, which clearly highlights the chromatin evidence of lineage-priming events before gene expression activation. Altogether, we demonstrate that analyses of chromatin accessibility mediated lineage-priming enable insight into the chromatin potential of cells and predict lineage fate outcomes.

## Discussion

High resolution, massively parallel simultaneous measurement of chromatin landscapes and gene expression in diverse tissues including during differentiation provided four key insights: (1) There is a high degree of congruence in the definition of differentiated cell types by both measures; (2) co-variation of chromatin and RNA across cells—within and between cell types—associates regulatory regions to their target genes; (3) among these, we identify DORCs, which reflect regulatory regions that control key lineage genes; and (4) focusing on DORCs, we find that chromatin activates prior to gene expression during differentiation, with chromatin potential foreshadowing RNA states of cells at longer time scales than RNA velocity. Furthermore, by downsampling reads or cells, we show these insights required the improved data quality and throughput (up to 10^6^ cells) of SHARE-seq.

To determine congruence, we find that both datasets largely reflect similar clusters of cell types demonstrating that cell types in tissues largely coordinate chromatin structure with transcription. Using this approach, we show that the joint data in SHARE-seq can provide excellent training for algorithms that aim to computationally map chromatin and RNA modalities across cells. Nevertheless, some cell states were not reflected equally in both profiles. In one example, a proliferative basal cell population was distinguished specifically in the gene expression dataset. In contrast, chromatin accessibility better distinguished cellular diversity within the TAC progenitor population, including chromatin-based signatures of lineage-priming.

To infer transcriptional regulation and recover key regulatory regions in differentiation, SHARE-seq provides a means to infer DORCs reflecting key lineage-determining genes. Leveraging SHARE-seq, it should now be possible to identify key regulatory regions, including developmental super-enhancers, and their associated target genes, without the need for isolating specific cell subsets or ChIP-seq experiments, which can be challenging for *in vivo* samples. We expect the inclusion of more layers of measurements, and improved computational methods for illustrating the differences between chromatin regulators and gene expression, will enable a more robust approach for defining chromatin-gene dynamics within complex tissues. This would be important in developmental biology, cancer research, and especially human genetics, where genetic variants associated with complex human diseases are found in non-coding regions, and relating them to specific cell types and target genes can be challenging.

Focusing on the incongruence between chromatin accessibility and gene expression, we demonstrate the existence of chromatin accessibility mediated lineage-priming, and define chromatin potential to describe the time difference upon hair follicle differentiation. Recently, RNA velocity approaches have predicted a cell’s future state from the difference between mature and nascent RNA. SHARE-seq allows us to make stronger predictions on a cell’s future potential in several ways. First, when we calculate chromatin potential, we relate the chromatin signal of one cell (or neighborhood) to the RNA signal in any cell (or neighborhood) from the same experiment, and can transverse longer time scales and identify cell fates earlier in differentiation. Second, leveraging the joint measurements of RNA (nascent and mature) and chromatin in every single cell, we can relate its current chromatin state to its current and future (by RNA velocity) states, to understand the distinction between its realized (in RNA) and as-yet-unrealized potential.

SHARE-seq provides a generalizable platform and opportunity to include additional layers of information per cell. With further development, we expect to integrate other scRNA-seq compatible measurements (Stuart et al., 2019), such as protein measurements (Stoeckius et al., 2017), genotyping, and lineage barcoding. Furthermore, powered by the massive scalability of this approach, SHARE-seq may be adapted for identifying RNA barcodes, particularly useful for CRISPR-based perturbation screens (Dixit et al., 2016). SHARE-seq may be further extended by replacing ATAC-seq with whole-genome transposition (Vitak et al., 2017) enabling methods for DNA methylation and chromatin conformation. In these efforts, scRNA-seq data may be used as a common scaffold for integration, providing a unique opportunity to comprehensively map between multiple layers of gene regulation, as well as to train algorithms that learn to map between different data modalities in a cell. As such, as we move toward a cell atlas, we anticipate SHARE-seq will likely play a key role in determining the full diversity of cell types and cell states, and the regulators that define them.

## Acknowledgements

We thank members of the Regev and Buenrostro lab for their critical reading of the manuscript and helpful discussions, including Fabiana Duarte, Inbal Avraham-Davidi, Bo Li, Eugene Drokhlyansky, and Karin Pelka. We are grateful to Jonathan Strecker in the Zhang lab for providing Tn5. J.D.B. and the Buenrostro lab acknowledge support by the Allen Distinguished Investigator Program through the Paul G. Allen Frontiers Group, the Chan Zuckerberg Initiative and support from the NIH New Innovator Award (DP2). A.R. in an Investigator of the Howard Hughes Medical Institute. Work was supported by the NHGRI Center for Cell Circuits (A.R.), the Klarman Cell Observatory (A.R.), a grant from the BRAIN initiative (A.R.), Smith Family Foundation Odyssey Award (Y.-C.H.), and NIH R01-AR070825 (Y.-C.H.). Y.-C.H. is a Pew Scholar and a NYSCF – Robertson Investigator. B.Z. is an awardee of the Charles A. King Trust Postdoctoral Research Fellowship.

## Author contributions

S.M. designed and optimized the protocol and generated datasets. B.Z. performed skin tissue dissociation and immunostaining. L.L. collected mouse lung tissue. Z.C., Y.H., V.K.K. C.L., J.D. provided computational support. A.B., T.L., Y.H. helped with experimental design. S.M., J.D.B. conducted the data analysis with input and guidance from A.R. J.D.B. and A.R. supervised the research. J.D.B., A.R., S.M. wrote the manuscript. All authors proofread the manuscript.

## STAR Methods

### Experimental methods

#### Mice

Mice were maintained in an Association for Assessment and Accreditation of Laboratory Animal Care (AAALAC) approved animal facility at Harvard University and MIT. Procedures were approved by the Institutional Animal Care and Use Committee of all institutions (institutional animal welfare assurance no. A-3125-01, 14-03-194 and 14-07-209).

#### Cell culture and tissue processing

##### (1) Cell culture

GM12878 cells were cultured in RPMI 1640 medium (11875-093, ThermoFisher) supplemented with 15% FBS (16000044, ThermoFisher) and 1% penicillin-streptomycin (15140122, ThermoFisher). NIH/3T3 and RAW 264.7 cells were cultured in Dulbecco’s Modified Eagle Medium (DMEM, 11965092, ThermoFisher) with the addition of 10% FBS and 1% of penicillin-streptomycin. Cells were incubated at 37°C in 5% CO_2_ and maintained at the exponential phase. NIH/3T3 and RAW 264.7 cells were digested with accutase for preparing single-cell suspension.

##### (2) Mouse skin

Female C57BL/6J mouse dorsal skins were collected at late anagen (P32). The hair cycle stages were confirmed using cryosectioning. To generate whole skin a single cell suspension, skin samples were incubated in 0.25% collagenase in HBSS at 37°C for 35-45 minutes on an orbital shaker. Samples were gently scraped from the dermal side and the single-cell suspension was collected by filtering through a 70µm filter followed by a 40µm filter. The epidermal portion of the skin samples were incubated in 0.25% trypsin-EDTA at 37°C for 35-45 minutes on the shaker and cells were gently scraped from the epidermal side. Single-cell suspensions were combined and centrifuged for 5 minutes at 4°C, resuspended in 0.25% FBS in PBS, and stained with DAPI (0.05µg/mL). Live cells were enriched by FACS. To enrich epidermal populations, CD140a negative population were purified by FACS and combined with whole skin cells in a ratio of 1:1.

##### (3) Mouse brain

An adult mouse brain was dissected, snap-frozen on dry ice, and stored at -80°C. A single nucleus suspension was prepared following the OMNI-ATAC protocol (Corces et al., 2017). Nuclei were resuspended in PBSI (0.1U/µl Enzymatics RNase Inhibitor, Y9240L, Qiagen; 0.05U/µl SUPERase inhibitor, AM2696, ThermoFisher; 0.04% Bovine Serum Albumin, BSA, 15260037, ThermoFisher in PBS).

##### (4) Mouse lung

Mouse lung was dissociated with fine scissors followed by proteolytic digestion using the Lung Dissociation kit (Miltenyi Biotech) following the manufacturer’s instructions. Dissociated cells were then incubated at 37°C for 20 minutes with rotation, then filtered using a 100µm strainer. Red blood cells were lysed using ACK buffer (A1049201, ThermoFisher).

#### Skin histology and immunofluorescence

Mouse skin samples were fixed in 4% paraformaldehyde (PFA) for 15 minutes at room temperature and then washed 6 times using PBS. The samples were immersed in 30% sucrose in PBS overnight at 4°C. Samples were cut and embedded in OCT (Sakura Finetek) and 35µM sections were harvested on positively charged slides. For immunohistochemistry, sections were fixed in 4% PFA for 2 minutes, washed with PBS and PBST. Sections were blocked with a blocking buffer (5% donkey serum, 1% BSA, 2% cold water fish gelatin, 0.3% Triton X-100 in PBS) for 1 hour at room temperature. Primary antibodies (anti-PolII S5, Abcam, ab5131; anti-PolII S2, Abcam, ab5095; anti-PolII, Abcam, ab817) were added and incubated overnight at 4°C. Secondary antibodies (anti-Rabbit IgG Alexa 488, Jackson ImmunoResearch, 711-545-152; anti-Mouse IgG Alexa 488, Jackson ImmunoResearch, 715-545-150) were added and incubated for 4 hours at room temperature.

#### SHARE-seq

##### (1) Preparing oligonucleotides for ligations

There are three barcoding rounds of hybridization reactions in SHARE-seq, with a different 96-well barcoding plate for each round (**Table S1**). Hybridization oligos have a universal linker sequence that is partially complementary to well-specific barcode sequences. These strands were annealed prior to cellular barcoding to create a DNA molecule with three distinct functional domains: a 5’ overhang that is complementary to the 3’ overhang present on the cDNA molecule or transposed DNA molecules (may originate from RT primer, transposition adapter or previous round of barcoding), a unique well-specific barcode sequence, and a 3’ overhang complementary to the 5’ overhang present on the DNA molecule to be subsequently ligated. Linker strands and barcode strands for the hybridization rounds were added to RNase-free 96-well plates to a total volume of 10µl/well with the following concentrations: round 1 plates contain 9µM round 2 linker strand and 10µM barcodes, round 2 plates contain 11µM round 2 linker strand and 12µM barcodes, and round 3 plates contain 13µM round 3 linker strand and 14µM barcodes. The oligos are dissolved in STE buffer (10 mM Tris pH 8.0, 50 mM NaCl, and 1 mM EDTA). Oligos are annealed by heating plates to 95°C for 2 minutes and cooling down to 20°C at a rate of -1°C per minute.

Blocking strands are complementary to the 3’ overhang present on the DNA barcodes used during hybridization barcoding rounds. Blocking occurs after well-specific barcodes have hybridized to cDNA molecules, but before all cells are pooled back together. The blocking step minimizes the possibility that unbound DNA barcodes mislabel cells in future barcoding rounds. 10µl of each blocking strand solution was added to each of the 96 wells after the first, second, and third round of hybridization of DNA barcodes, respectively. Blocking strand solutions were prepared at a concentration of 22µM for round 1, 26.4µM for round 2, and 23µM for round 3. Blocking strands for the first two rounds were in a 2× T4 DNA Ligase buffer (NEB) while the third round was in 0.1% Triton X-100. Both ligation reaction and blocking reaction were incubated with cells for 30 minutes at room temperature with gentle shaking (300 rpm). All the oligos are thawed to room temperature before using.

##### (2) Fixation

For simplicity, cells and nuclei, which were processed identically for the following steps, are both referred to as cells. Cells were centrifuged at 300g for 5 minutes and resuspended to 1 million cells/ml in PBSI. Cells were fixed by adding formaldehyde (28906, ThermoFisher, final concentration of 0.1% for cell lines or 0.2% for primary tissues) and incubated at room temperature for 5 minutes. The fixation was stopped by adding 56.1µl of 2.5M glycine, 50µl of 1M Tris-HCl pH 8.0, and 13.3 µl of 7.5% BSA on ice. The sample was incubated at room temperature for 5 minutes and then centrifuged at 500g for 5 minutes to remove supernatant. All centrifugations were performed on a swing bucket centrifuge. The cell pellet was washed twice with 1ml of PBSI, and centrifuged at 500g for 5 minutes between washings. The cells were resuspended in PBS with 0.1U/µl Enzymatics RNase Inhibitor and aliquoted for transposition.

##### (3) Transposition

The transposition reaction is performed similarly to previous published work (Corces et al., 2017) with minor modifications. All the oligos used in this protocol can be found in **Table S1**. The 100µM Read1 and phosphorylated Read2 oligos were annealed with an equal amount of 100µM blocked ME-complement oligo by heating at 85°C for 2 minutes and slowly cooling down to 20°C at a ramp rate of -1°C/minute. The annealed oligos were mixed with an equal volume of cold glycerol and stored at -80°C until use. In-house produced Tn5^10^ was mixed with an equal volume of dilution buffer (50 mM Tris, 100 mM NaCl, 0.1 mM EDTA, 1 mM DTT, 0.1% NP-40, and 50% glycerol). Diluted Tn5 was then mixed with an equal volume of annealed oligos and incubated at room temperature for 30 minutes before transposition.

For each transposition reaction, cells (10,000-20,000 cells in 5µl PBSI) and 42.5µl of transposition buffer (38.8 mM Tris-acetate, 77.6 mM K-acetate, 11.8 mM Mg-acetate, 18.8% DMF, 0.12% NP-40, 0.47% Protease Inhibitor Cocktail, and 0.8 U/µl Enzymatics RNase Inhibitor) were mixed and incubated at room temperature for 10 minutes. 2.5µl of assembled Tn5 was added to the transposition reaction. Depending on the target number of cells to be recovered, the number of transposition reactions can be scaled up. In general, we prepare 10-40 reactions, which is equivalent to 100,000-800,000 cells. The transposition was carried out at 37°C for 30 minutes with shaking at 500rpm. The sample was centrifuged at 1,000g for 3 minutes and then washed with 1ml Nuclei Isolation Buffer (NIB) (10mM Tris buffer pH 7.5, 10mM NaCl, 3mM MgCl_2_, 0.1% NP-40, 0.1U/µl Enzymatics RNase Inhibitor, and 0.05U/µl SUPERase RI). The sample was then resuspended to 60µl of NIB and before proceeding to reverse transcription.

##### (4) Reverse transcription

Transposed cells (60µl) were mixed with 240µl of RT mix (1.25× RT buffer, 0.5 U/µl Enzymatics RNase Inhibitor, 625µM dNTP, 12.5µM RT primer with an affinity tag, 18.75% PEG 6000, and 25 U/µl Maxima H Minus Reverse Transcriptase). The RT primer contains a poly-T tail, a Unique Molecular Identifier (UMI), a universal ligation overhang, and a biotin molecule. The sample was heated at 50°C for 10 minutes, then went through 3 thermal cycles (8°C for 12s, 15°C for 45s, 20°C for 45s, 30°C for 30s, 42°C for 120s and 50°C for 180s), and finally incubated at 50°C for 5 minutes. After reverse transcription, 300µl of NIB was added and the sample was centrifuged at 1,000g for 3 minutes to remove supernatant. Cell pellet was washed with 0.5ml of NIB and centrifuged at 1,000g for 3 minutes. Cells were resuspended in 4,608µl of hybridization mix (1× T4 ligation buffer, 0.32 U/µl Enzymatics RNase Inhibitor, 0.05 U/µl SUPERase RI, 0.1% Triton X-100, and 0.25× NIB).

##### (5) Hybridization and ligation

Cells in ligation mix (40µl) were added to each of the 96 wells in the first-round barcoding plate. Each well already contained 10µl of the appropriate DNA barcodes. The round 1 barcoding plate was incubated for 30 minutes at room temperature with gentle shaking (300 rpm) to allow hybridization to occur before adding blocking strands. 10µl of round 1 blocking oligo was added and the plate was incubated for 30 minutes at room temperature with gentle shaking (300 rpm). Cells from all 96 wells were combined into a single multichannel basin. Subsequent steps in round 2 and round 3 were identical to round 1, except that 50µl and 60µl of pooled cells were split and added to barcodes in round 2 (total volume of 60µl/well) and round 3 (total volume of 70µl/well), respectively. After adding the round 3 blocking oligo, cells from all wells were combined and centrifuged at 1,000g for 3 minutes to remove supernatant. The cell pellet was washed twice with 1ml of NIB, and centrifuged at 1,000g for 3 minutes between washings. Cells were re-suspended in the ligation mix (1× T4 ligation buffer, 0.32 U/µl Enzymatics RNase Inhibitor, 20 U/µl T4 DNA ligase (M0202L, NEB), 0.1% Triton X-100, 0.2× NIB) and incubated for 30 minutes at room temperature with gentle shaking (300 rpm). Cells were washed once with 0.5ml washing buffer and resuspended in 100µl of NIB, counted and aliquoted to 0.2ml PCR tubes with 1,000-20,000 cells per tube.

##### (6) Reverse crosslinking and affinity pull-down

NIB was added to each sample to bring the volume to 50µl in total. 50µl of 2× reverse crosslinking buffer (100mM Tris pH 8.0, 100mM NaCl, and 0.04% SDS), 2µl of 20mg/ml proteinase K, and 1µl of SUPERase RI were mixed with each sample and incubated at 55°C for 1 hour. 5µl of 100mM PMSF was added to the reverse crosslinked sample to inactivate proteinase K and incubated at room temperature for 10 minutes. For each sample, 10µl of MyOne C1 Dynabeads were washed twice with 1× B&W-T buffer (5mM Tris pH 8.0, 1M NaCl, 0.5mM EDTA, and 0.05% Tween 20) and once with 1× B&W-T buffer supplemented with 2U/µl SUPERase RI. After washing, the beads were resuspended in 100µl of 2× B&W buffer (10mM Tris pH 8.0, 2M NaCl, 1mM EDTA, and 4U/µl SUPERase RI) and mixed with the sample. The mixture was rotated on an end-to-end rotator at 10 rpm for 60 minutes at room temperature. The lysate was put on a magnetic stand to separate supernatant and beads.

##### (7) scATAC-seq library preparation

The supernatant that contained the transposed DNA fragments was purified with DNA clean and concentrator kit and eluted to 10µl of Tris buffer (pH 8.0). Fragments were PCR amplified with Ad1 primer with sample barcodes and P7 primer. The amplification procedure was similar to standard bulk ATAC-seq library preparation (Corces et al., 2017) with minor modifications: the annealing temperature was set to 65°C instead of 72°C.

##### (8) cDNA library preparation

Beads were washed three times with 1× B&W-T buffer and once with STE (10mM Tris pH 8.0, 50mM NaCl, and 1mM EDTA) both supplemented with 1U/µl SUPERase inhibitor. Beads were resuspended in 50µl of template switch mix (15% PEG 6000, 1× Maxima RT buffer, 4% Ficoll PM-400, 1mM dNTPs, 1U/µl Enzymatics RNase-In, 2.5µM TSO, and 10U/µl Maxima H Minus Reverse Transcriptase). Beads were rotated on an end-to-end rotator at 10rpm for 30 minutes at room temperature, and then shaken at 300rpm for 90 minutes at 42°C. Beads were resuspended by pipetting every 30 minutes during agitation. After template switching, 100µl of STE were added to each tube to dilute the sample. The supernatant was removed by placing the sample on a magnetic stand. Beads were washed with 200µl of STE without disturbing the bead pellet. Beads were then resuspended in 55µl of PCR mix (1× Kapa HiFi PCR mix, 400nM P7 primer, and 400nM RNA PCR primer). The PCR reaction was carried out at the following conditions: 95°C for 3 minutes, and then thermocycling 14 cycles at 98°C for 30s, 65°C for 45s and 72°C for 3 minutes. Optionally, we run 5 cycles of PCR, take a 2.5µl sample, added 7.5µl of PCR cocktail with 1× EvaGreen (Biotium), and run qPCR. The qPCR reactions were amplified to saturation to determine the number of cycles required for the remaining samples on the plate. The qPCR reaction was carried out at the following conditions: 95°C for 3 minutes, and then 20 thermal cycles at 98°C for 30s, 65°C for 20s and 72°C for 3 minutes. Libraries were amplified for 12-14 cycles in total for 1,000 cells. Amplified cDNA was purified by 0.8× (for cell line) or 0.6× (for primary tissue) AMPure beads and eluted to 10µl of Tris pH 8.0 buffer. The amount of cDNA was quantified by Qubit (ThermoFisher).

##### (9) Tagmentation and scRNA-seq library preparation

100µM Read1 oligo was annealed with an equal amount of 100µM blocked ME-complement oligo and assembled with Tn5 as described above. For each sample, 50ng cDNA was fragmented in a 50µl tagmentation mix (1× TD buffer from Illumina Nextera kit, and 5 µl assembled Tn5) at 55°C for 5 minutes. Fragmented cDNA was purified with the DNA Clean and Concentrator kit (Zymo) and eluted to 10µl of Tris pH 8.0 buffer. Purified cDNA was then mixed with tagmentation PCR mix (25µl of NEBNext High-Fidelity 2× PCR Master Mix, 1µl of 25µM P7 primer and 1µl of 25µM Ad1 primer with sample barcodes). PCR was carried out at the following conditions: 72°C for 5 minutes, 98°C for 30s, and then 7 cycles at 98°C for 10s, 65°C for 30s and 72°C for 1 minute. The amplified library was purified by 0.7× AMpure beads and eluted to 10µl of Tris buffer (pH 8.0).

##### (10) Quantification and sequencing

Both scATAC-seq and scRNA-seq libraries were quantified with KAPA Library Quantification Kit and pooled for sequencing. Libraries were sequenced on the Next-seq platform (Illumina) using a 150-cycle High-Output Kit (Read 1: 30 cycles, Index 1: 99 cycles, Index 2: 8 cycles, Read 2: 30 cycles) or the Nova-seq platform (Illumina) using a 200-cycle S1 kit (Read 1: 50 cycles, Index 1: 99 cycles, Index 2: 8 cycles, Read 2: 50 cycles).

### Computational methods

#### SHARE-ATAC-seq pre-processing

Raw sequencing reads were trimmed with a custom python script. Reads were aligned to hg19 or mm10 genome using bowtie2 (Langmead et al. 2012) with (-X2000) option. For each read, there are four sets of barcodes (eight bases each) in the indexing reads. The data were demultiplexed tolerating one mismatched base in each 8-base barcode. Reads with alignment quality < Q30, improperly paired, mapped to the unmapped contigs, chrY, and mitochondria, were discarded. Duplicates were removed using Picard tools (http://broadinstitute.github.io/picard/). Open chromatin regions peaks were called on individual samples using MACS2 peak caller (Zhang et al., 2008) with the following parameters: --nomodel –nolambda –keep-dup -call-summits. Peaks from all samples were merged and peaks overlapping with ENCODE blacklisted regions (https://sites.google.com/site/anshulkundaje/projects/blacklists) were filtered out. Peak summits were extended by 150bp on each side and defined as accessible regions. Peaks were annotated to genes using Homer (Heinz et al., 2010). The fragment counts in peaks and TF scores were calculated using chromVAR (Schep et al., 2017).

#### SHARE-RNA-seq pre-processing

Base calls were converted to the fastq format using bcl2fastq. Reads were trimmed with a custom python script. We removed reads that do not have TTTTTT at the beginning of Read 2 allowing one mismatch. Reads were aligned to the mouse genome (version mm10) using STAR (Dobin et al. 2013) (STAR --chimOutType WithinBAM --outFilterMultimapNmax 20 --outFilterMismatchNoverLmax 0.06 --limitOutSJcollapsed 2000000). For species mixing experiments, reads were aligned to a combined human (hg19) and mouse (mm10) genome and only primary alignments were considered. Data were demultiplexed tolerating one mismatched base in each 8-base barcode. Aligned reads were annotated to both exons and introns using featurecounts (Liao et al. 2014). To speed up processing, only barcode combinations with >100 reads were retained. UMI-Tools (Smith et al. 2017) was used to collapse UMIs of aligned reads that were within 1nt mismatch of another UMI. UMIs that were only associated with one read were removed as potential ambient RNA contamination. A matrix of gene counts by cell was created with UMI-Tools. For cell line data, cells that expressed >7,500 genes, <300 genes, or >1% mitochondrial reads were removed. For tissue data, cells that expressed >10,000 genes, <100 genes, or >2% mitochondrial reads were removed. Expression counts (number of transcripts) for a given gene in a given cell were determined by counting unique UMIs and compiling a Digital Gene Expression (DGE) matrix. Mitochondrial genes are removed. Seurat V3 (Stuart et al. 2019) was used to scale the DGE matrix by total UMI counts, multiplied by the mean number of transcripts, and values were log transformed. To visualize data, the top 3,000 variable genes were projected into 2D space by UMAP (McInnes et al. 2018). Ambient RNA level was estimated using a previously reported approach (Ding et al., 2019).

#### Peak-gene *cis*-association and DORC identification

To calculate peak-gene associations in *cis*, we considered all ATAC peaks that are located in the ± 50 kb or ± 500 kb window around each annotated TSS. We used peak counts and gene expression values to calculate the observed Spearman correlation (obs) of each peak-gene pair. To estimate the background, we used chromVAR to generate 100 background peaks for each peak by matching accessibility and GC content, and calculated the Spearman correlation coefficient between those background peaks and the gene, resulting in a null peak-gene Spearman correlation distribution that is independent of peak-gene proximity. We calculated the expected population mean (pop.mean) and expected population standard deviation (pop.sd) from expected Spearman correlations. The Z score is calculated by z=(obs-pop.mean)/pop.sd, and converted to a *p*-value based on the normal distribution. For peaks associated with multiple genes, we only kept peak-gene associations with the smallest *p*-value.

To define DORCs (a set of nearby peaks per gene), we rank genes by the number of significantly associated peaks (± 50 kb around TSSs, *p* < 0.05). We used 10 and 5 peaks per gene as cutoffs for skin data and GM12878 data, respectively. We then re-calculate peak-gene association by expanding the window to ±500 kb around TSSs. The DORC score was calculated by summing up all the significantly correlated peak counts per gene, and then normalized by dividing the total unique fragments in peaks.

#### TF-gene correlation in *trans*

We used TF scores derived from chromVAR and gene expression values to calculate the observed Spearman correlation (obs) of each TF-gene pair. TF scores were root-mean-square normalized and gene expression values were normalized using the SCtransform function in Seurat. Z scores and *p*-values were calculated in the same way in the *cis-*analysis.

#### Comparison to other technologies

We compared the performance of SHARE-seq to sci-CAR (Cao et al., 2018), SNARE-seq (Chen et al., 2019) and Paired-seq (Zhu et al., 2019) using cell line data. We used deeply sequenced GM12878 data for SHARE-seq, published A549 cell line data for sci-CAR (Cao et al., 2018) and published cell line mixture data for SNARE-seq (Chen et al., 2019) and Paired-seq (Zhu et al., 2019). We used the authors’ count matrices, which were obtained on libraries that were sequenced to saturation. For each assay, we determine the cutoff by ranking the number of unique molecules per cell barcode. We set cutoff at the steep drop-off which indicates separation between the cell-associated barcodes and the barcodes associated with debris (**Supplementary note Fig. 1**).

To compare SHARE-seq with other high-throughput scATAC-seq methods using cell line data, we used the approach described in previous paper (Lareau et al., 2019), and compared with published datasets, including Cusanovich et al.(Cusanovich et al., 2015) (GSE67446), Pliner et al. (Pliner et al., 2018) (GSE109828), Preissl et al. (Preissl et al., 2018) (GSE1000333), Lareau et al. (Lareau et al., 2019) (GSE123581), and Buenrostro et al. (Buenrostro et al., 2015) (GSE65360).

To compare scATAC-seq technologies in primary tissue, we generated sci-ATAC, SHARE-seq, and 10x Genomics scATAC-seq datasets on adult mouse lung using the same sample processing method (above).

To compare SHARE-seq with other high-throughput scRNA-seq/snRNA-seq methods, we processed four adult mouse brain datasets the same way as SHARE-seq. We downloaded count matrix for nuclei (https://support.10xgenomics.com/single-cell-gene-expression/datasets/2.1.0/nuclei_2k) and cells (Zeisel et al., 2018) processed by 10x Genomics (P60 cortex, SRP135960), cells processed by Drop-seq (Saunders et al., 2018) (P60 Cortex, GSE116470), and nuclei processed by DroNc-seq (Habib et al., 2017) (PFC, GSE71585).

#### Cell cycle signature

To calculate the cell cycle signature, we used our previously published cell cycle gene list (Tirosh et al. 2016) and summed up the normalized cell cycle gene counts per cell. We did not regress out the cell cycle signature, because it is one of the most important signatures in TACs.

#### Computational pairing

To confirm if computational pairing methods correctly predict cell type in scATAC-seq based on a scRNA-seq profile, we used Seurat v3.0 (Stuart et al. 2019) to calculate gene activity scores from scATAC-seq. Next, we identified anchors between the scATAC-seq and scRNA-seq datasets using CCA (Stuart et al. 2019) and used these anchors to transfer cell-type labels from scRNA-seq to scATAC-seq. We calculated the percent of mismatch between the predicted cell type to the actual cell type.

#### Brain data analysis

For the brain sample, we aggregated scATAC-seq data generated using SureCell (Lareau et al., 2019) as pseudo-bulk samples, then extracted a small number of principal components (PCs) from the normalized pseudo-bulk count matrix. We next projected the scATAC-seq data to the space spanned by the PCs. The projected data was then visualized using tSNE and UMAP. To jointly cluster on ATAC and RNA signal, we used Similarity NEtwork FUSION (Wang et al. 2014) to combine the distance matrix in chromatin space and RNA space. After generating the fused distance matrix, we then calculated *k*-nearest neighbor graph and found clusters using the Louvain community detection algorithm. The clusters were assigned based on both marker gene and scATAC-seq signal.

#### Skin scATAC-seq peak count matrix

To ensure our peak set in skin includes ATAC peaks from rare populations, we performed two rounds of peak calling. We first called peaks on filtered reads from all cells and generated 1^st^-round cell-peak count matrix. We then filtered cells based on both ATAC and RNA profiles and identified clusters based on RNA profiles. We next called peaks again on aggregated pseudo bulk samples from each cluster and merged all peak summits, to generate a 2^nd^-round cell-peak count matrix.

#### Skin scATAC-seq dimension reduction

To reduce the dimension of ATAC-seq data, we tested cisTopic (González-Blas et al. 2019), chromVAR motif score and Kmer (Schep et al. 2017), and snapATAC (Fang et al. 2019) approaches.

#### Pseudotime inference

To calculate pseudotime based on scATAC-seq data of TACs, IRS and Hair Shaft populations, we provided 55 normalized topics from cisTopic as input to Palantir (Setty et al. 2019). We then defined lineages based on the probability of lineage assignment.

#### Residual analysis

Both DORC scores and gene expression were smoothed over pseudotime with local polynomial regression fitting (loess) separately, then min-max normalized. The residual for each gene was calculated by subtracting normalized gene expression from normalized DORC scores.

#### Chromatin potential

To calculate chromatin potential, we first smoothed DORC scores (chromatin space) and corresponding gene expression (RNA space) over a *k*-nearest neighbor graph (*k*-NN, *k* = 50), calculated using normalized ATAC topics from cisTopic. Next, we calculated another *k*-NN (*k* = 10), between the smoothed chromatin profile of a given cell (C_atac, i_), and the smoothed gene expression profile of each cell (C_rna, i, j_). We then calculated the distance (D_i, j_) between the C_atac, i_ and the average of C_rna, j_ in chromatin space. The arrow length is defined by normalizing D_i, j_. For visualization, we smoothed arrows with the 15 *k*-NNs in low dimensional space. For grid view, we divided the UMAP space into 40 × 40 grid, then averaged the arrows for all the cells within each grid.

#### RNA velocity

RNA velocity was calculated using Velocyto (La Manno et al. 2018) with default settings. For visualization, we smoothed arrows with the 15 RNA *k*-NNs. For grid view, we divided the UMAP space into 40 × 40 grid, then averaged the arrows for all the cells within each grid.

