## Supplementary file for "Chromatin potential identified by shared single cell profiling of RNA and chromatin"

### Supplemental Information

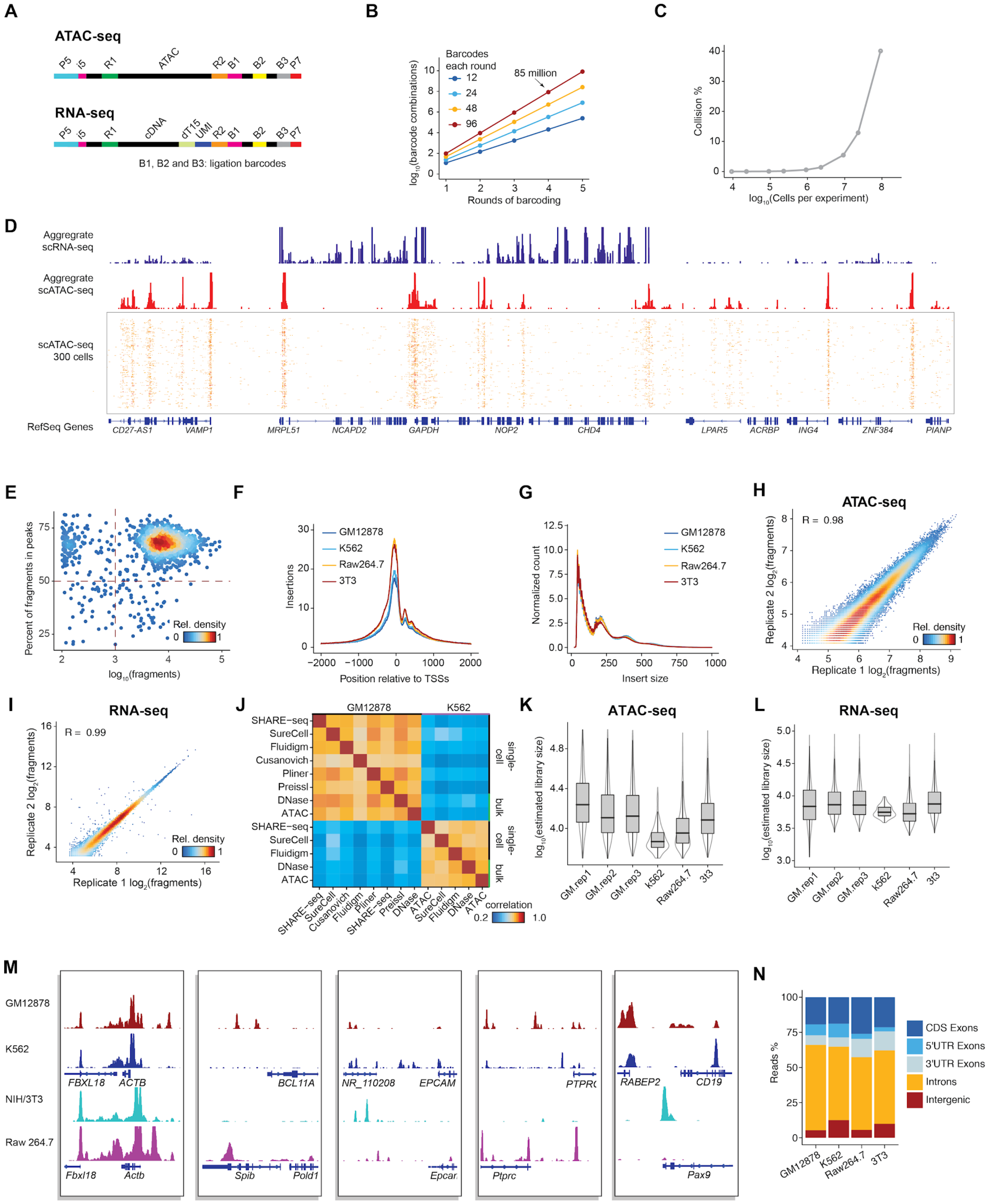


**Figure S1. The principle of SHARE-seq and data quality control on cell line datasets.**
(**A**) The structure of scATAC-seq and scRNA-seq sequencing library.
(B) The expected number of barcode combinations exponentially scales with the rounds of barcoding.

(C) Expected barcode collision happens with a large number of cells (> $10^5$).
(D) Aggregate single-cell accessibility and gene expression profiles in GM12878 cells.
(E) Scatter plot of the portion of reads in peaks (FRIP) of GM12878 ATAC-seq data.
(F) The enrichment of ATAC-seq reads around TSSs.
(G) The insert size distribution of ATAC-seq fragments.
(H,I) The SHARE-seq reproducibility between biological replicates on ATAC-seq (H) and RNA-seq (I).
(J) Aggregated ATAC-seq portion of SHARE-seq profile compares to Cusanovich (Cusanovich et al., 2015), Pliner (Pliner et al., 2018), Preissl (Preissl et al., 2018), SureCell (Lareau et al., 2019), sci-ATAC-seq (LaFave et al. under review), Flugidm C1 dataset (Buenrostro et al., 2015), and DNase-seq (ENCODE).
(K,L) The estimated library size (the unique molecules could be recovered by sequencing to saturation, estimated based on the duplication rate and recovered unique molecule) in SHARE-ATAC-seq (K) and SHARE-RNA-seq (L).
(M) The aggregated single-cell SHARE-seq accessibility profiles across different cell lines.
(N) The RNA reads distribution of SHARE-seq in the genome.

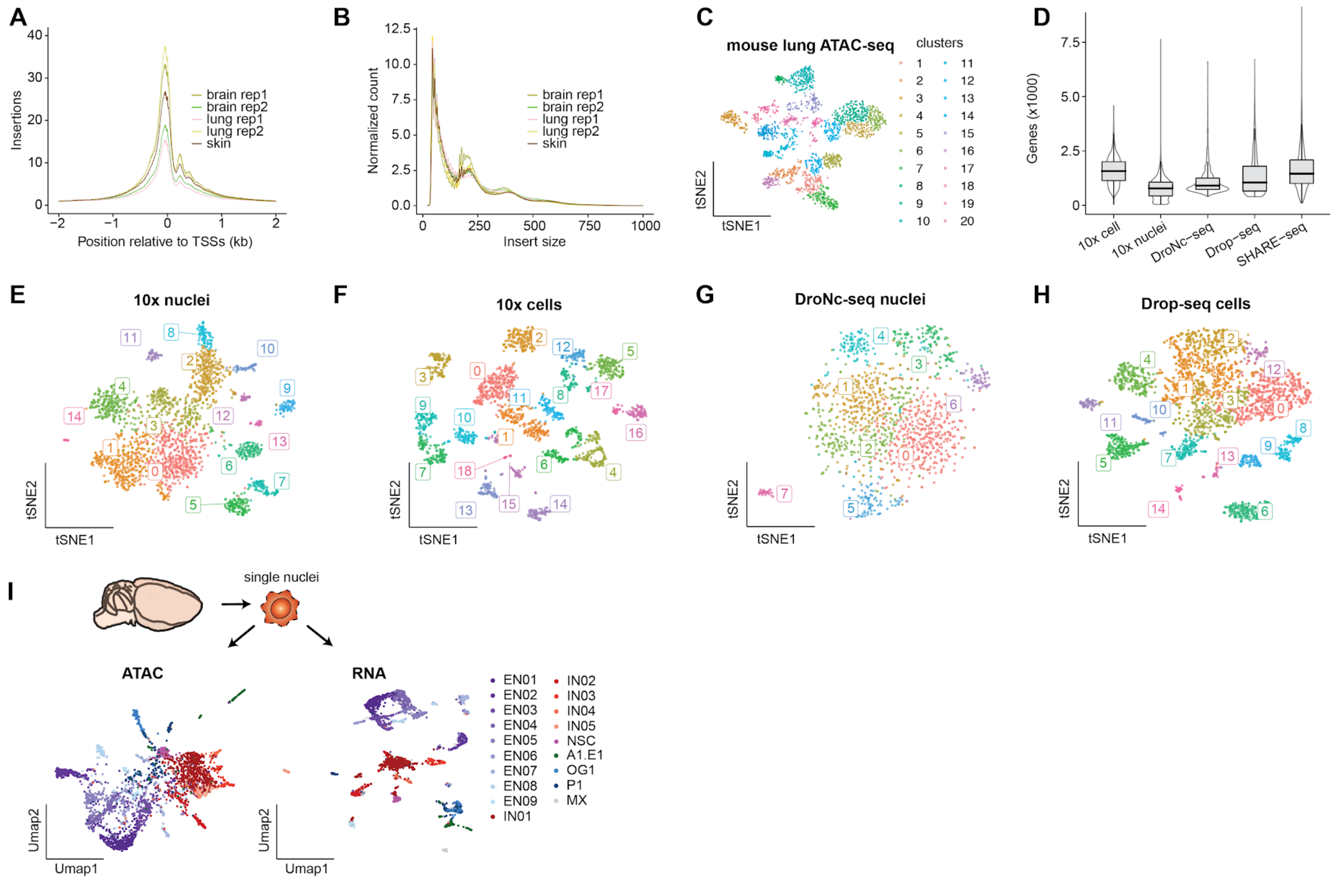


**Figure S2. Quality control for scATAC libraries on multiple tissues generated using SHARE-seq protocol.**
(A) The enrichment of ATAC-seq reads around TSSs.
(B) The insert size distribution of ATAC-seq fragments.
(C) t-SNE clustering of SHARE-ATAC-seq on adult mouse lung.
(D) Comparison of SHARE-RNA-seq to previously deposited 3' single cell/nuclei adult mouse brain datasets (**STAR Methods**) in terms of the number of genes detected.
(E-H) t-SNE visualizations of other 3' single cell/nuclei adult mouse brain dataset to compare to SHARE-RNA-seq. The numbers denote the clusters identified with the Louvain community detection algorithm.
(I) ATAC UMAP and RNA UMAP colored by the cell type assigned by the joint clustering of ATAC-seq and RNA-seq data in the mouse brain.

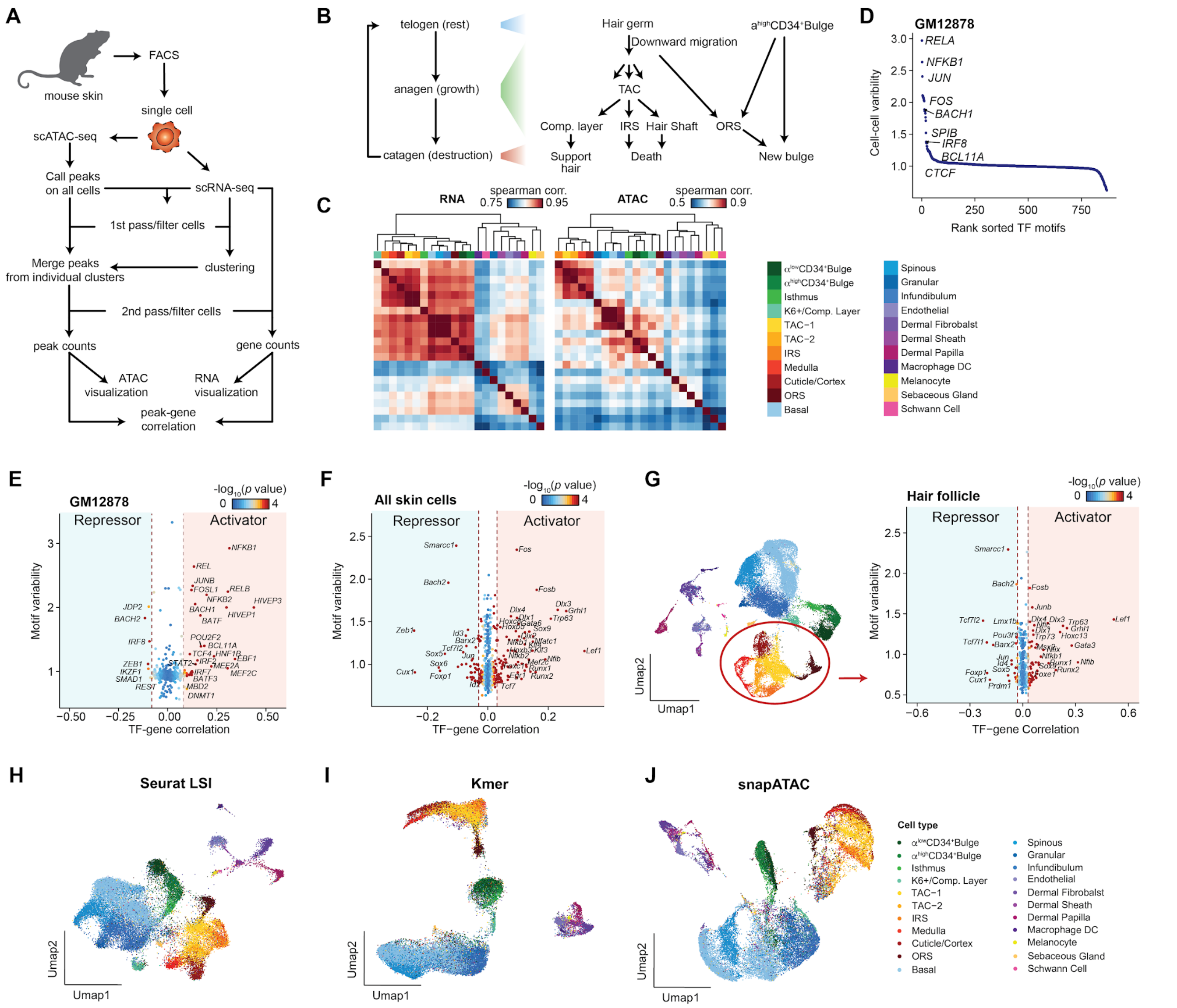


**Figure S3. SHARE-seq enables joint profiling of chromatin accessibility and transcription in adult mouse skin.**

(A) Schematic of a computational pipeline to process SHARE-seq data on adult mouse skin.

(B) The hair follicle cell types shift during hair follicle cycles.

(C) The pairwise similarity across RNA and ATAC clusters. Peak counts are used for computing correlation of the ATAC-seq data.

(D) The cell-cell variability of TF motif scores in the GM12878 cell line.

(E-G) The TF motif scores to gene expression correlation in GM12878 cells (E), all skin cells (F), and hair follicle cells (G). The dots color denotes the significance of the correlation.

(H-J) ATAC UMAP visualization with Seurat LSI (H), chromVAR Kmer (I), and snapATAC (J) approaches (**STAR Methods**). Points are colored by clusters labels.

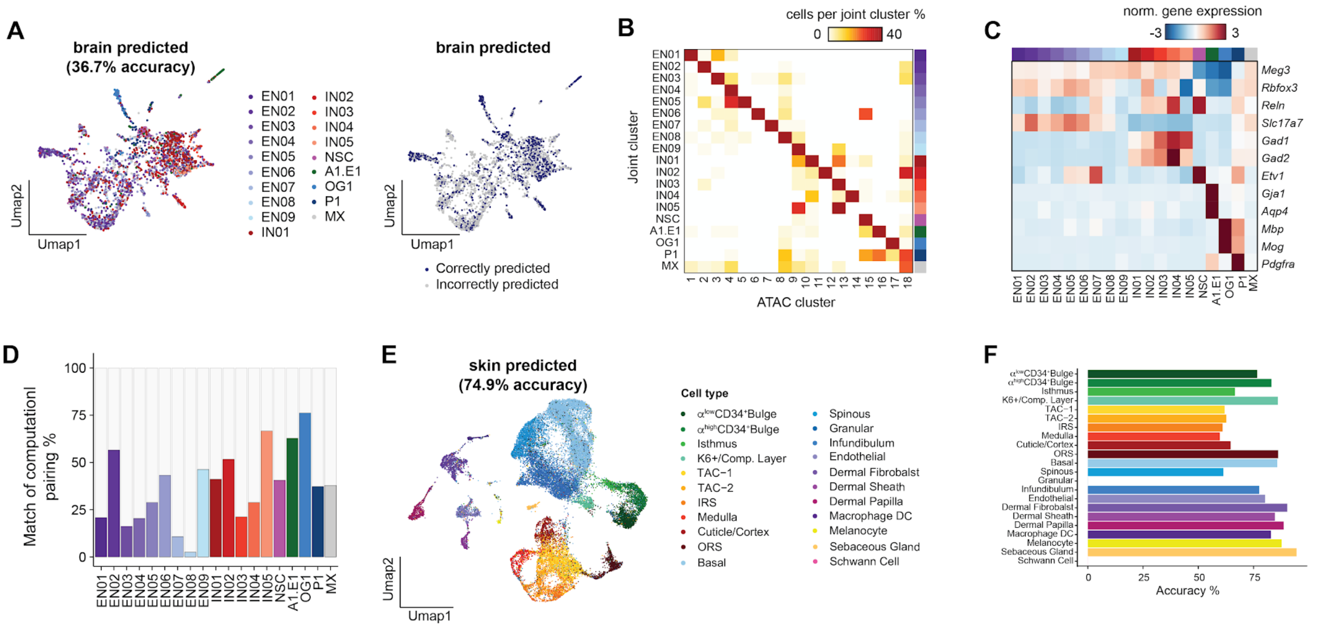


**Figure S4. The computational paring of ATAC-RNA misassigns cell types.**

(A) ATAC UMAP colored by computationally inferred cell type in the mouse brain. The computational pairing was performed by transferring the assigned cluster label to the ATAC cluster using Seurat (Stuart et al., 2019).

(B) Heatmap showing the proportion of cells in the joint cluster that overlaps in ATAC clusters in the mouse brain.

(C) Marker genes for each assigned cell type in the mouse brain.

(D) Histogram showing the percentage of cells that are correctly computationally assigned for each cell type in the mouse brain.

(E) UMAP visualization of computationally inferred cell type in mouse skin. The cell type labels are transferred from RNA-seq to ATAC-seq using Seurat (Stuart et al., 2019).

(F) Histogram showing the percentage of cells that are correctly computationally assigned for each cell type in mouse skin.

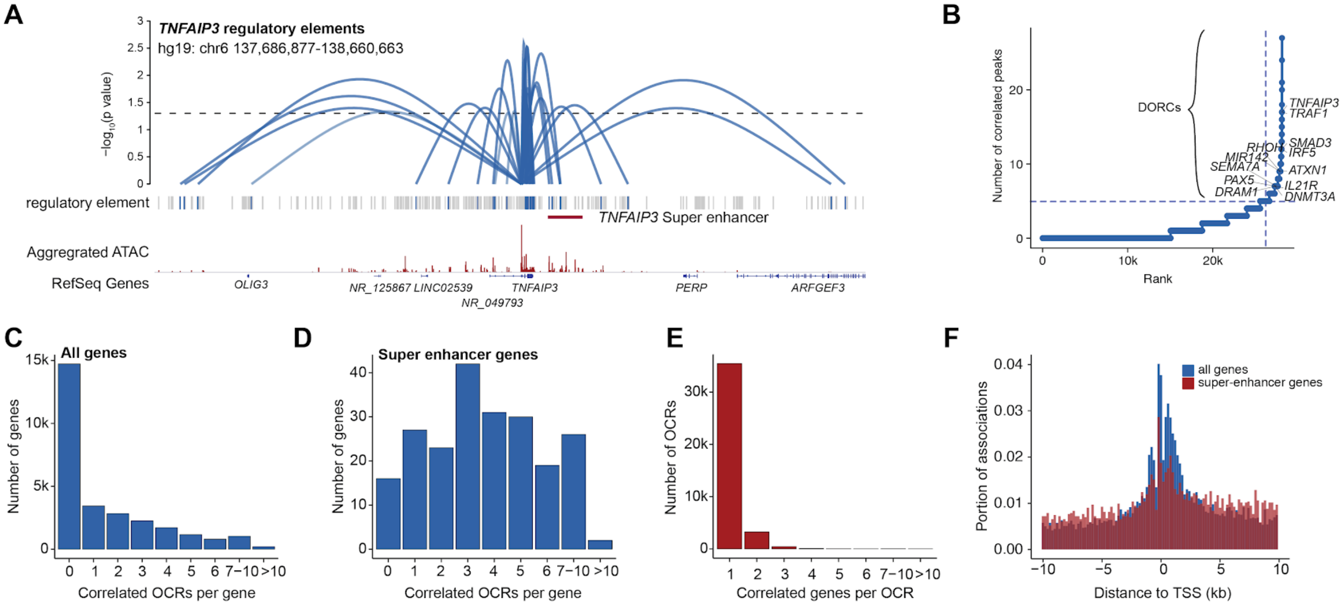


**Figure S5. SHARE-seq reveals *cis*-regulation within a cell line (GM12878).**
(A) Loops denote the correlation of peak accessibility and RNA expression at the *TNFAIP3* locus, loop height represents the significance of the correlation. Super-enhancer annotation generated from GM12878 cell line (Hnisz et al., 2013).
(B) The number of significant peak-gene associations for each gene.
(C,D) Histogram of the number of significant peak-gene associations per gene for all the genes (C) super-enhancer related genes (D).
(E) The number of genes associated with each significant peak.
(F) The distance of each significant peak-gene association ($p < 0.05$) to the TSS of each gene.

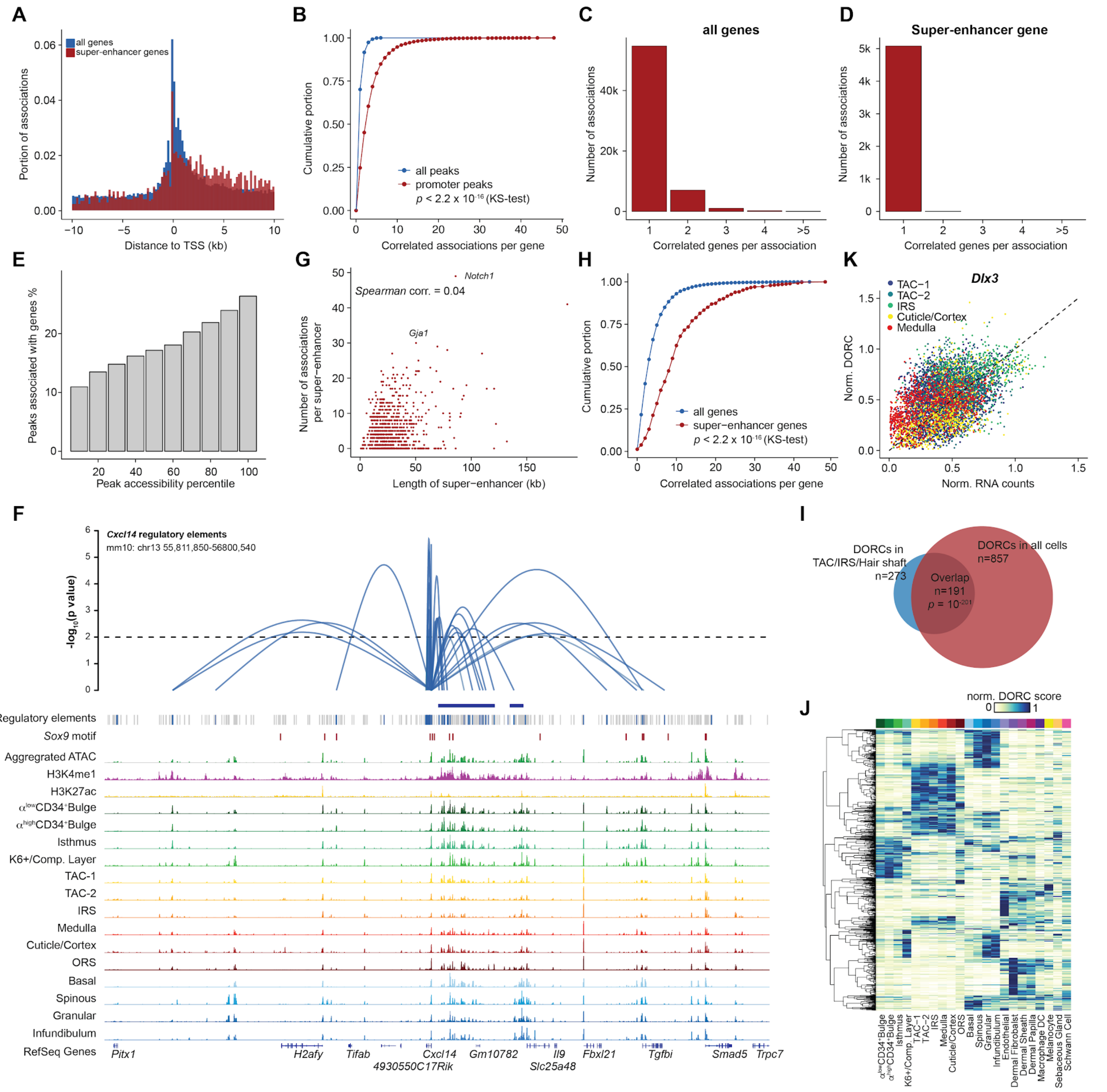

**Figure S6. *Cis*-regulation regions overlap with known super-enhancers, and are gene- and cell stage-specific.**

(A) Distance of each significant peak-gene association ($p < 0.05$) to the TSS of each gene.

(B) A cumulative distribution function plot of peak-gene associations across all peaks and promoter peaks.

(C,D) The number of genes associated with each significant peak for all genes (C) and super-enhancer related genes (D).

(E) The portion of peaks associated with genes varies with chromatin accessibility level.

(F) Loops denote the correlation of peak accessibility and RNA expression around the *Cxcl14* locus, loop height represents the significance of the correlation. H3K4me1 and H3K27ac ChIP-seq tracks and super-enhancer annotation generated from isolated TAC population (Adam et al., 2015).

(G) The scatter plot showing the length of super-enhancer is not correlated with the number of associated peaks.
(H) A cumulative distribution function plot of peak-gene associations for each gene.
(I) The overlapping between DORCs identified in TAC/IRS/Hair shaft and in all cells.
(J) DORC activity for each defined cluster, values are normalized by the min and max activity.
(K) Scatter plot of the *Dlx3* DORC score and *Dlx3* gene expression.

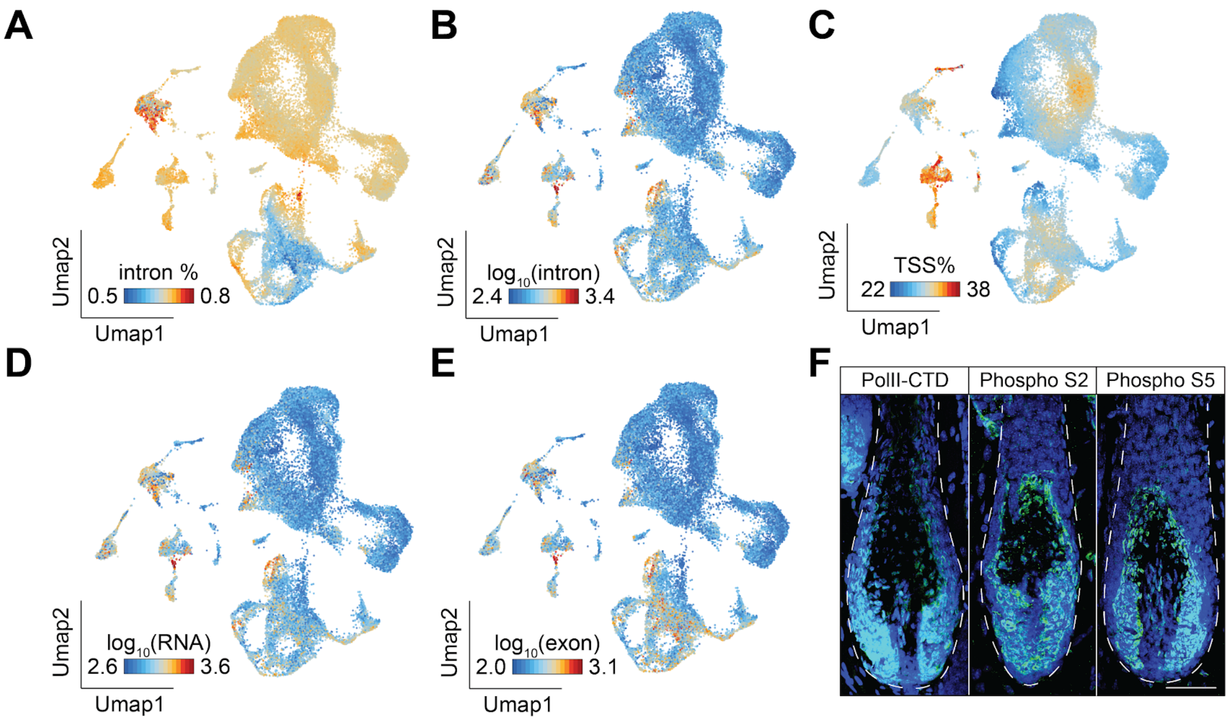


**Figure S7. TACs show a low percentage of RNA intronic reads.**
(A-E) RNA UMAP visualization of the percentage of intronic RNA-seq reads (A), total intronic RNA-seq reads (B), the percentage of promoter ATAC-seq reads (C), the yield of unique transcripts (D), and total exonic RNA-seq reads (E).
(F) Immuno-staining of PolII, PolII S2, and PolII S5 in the late anagen stage.

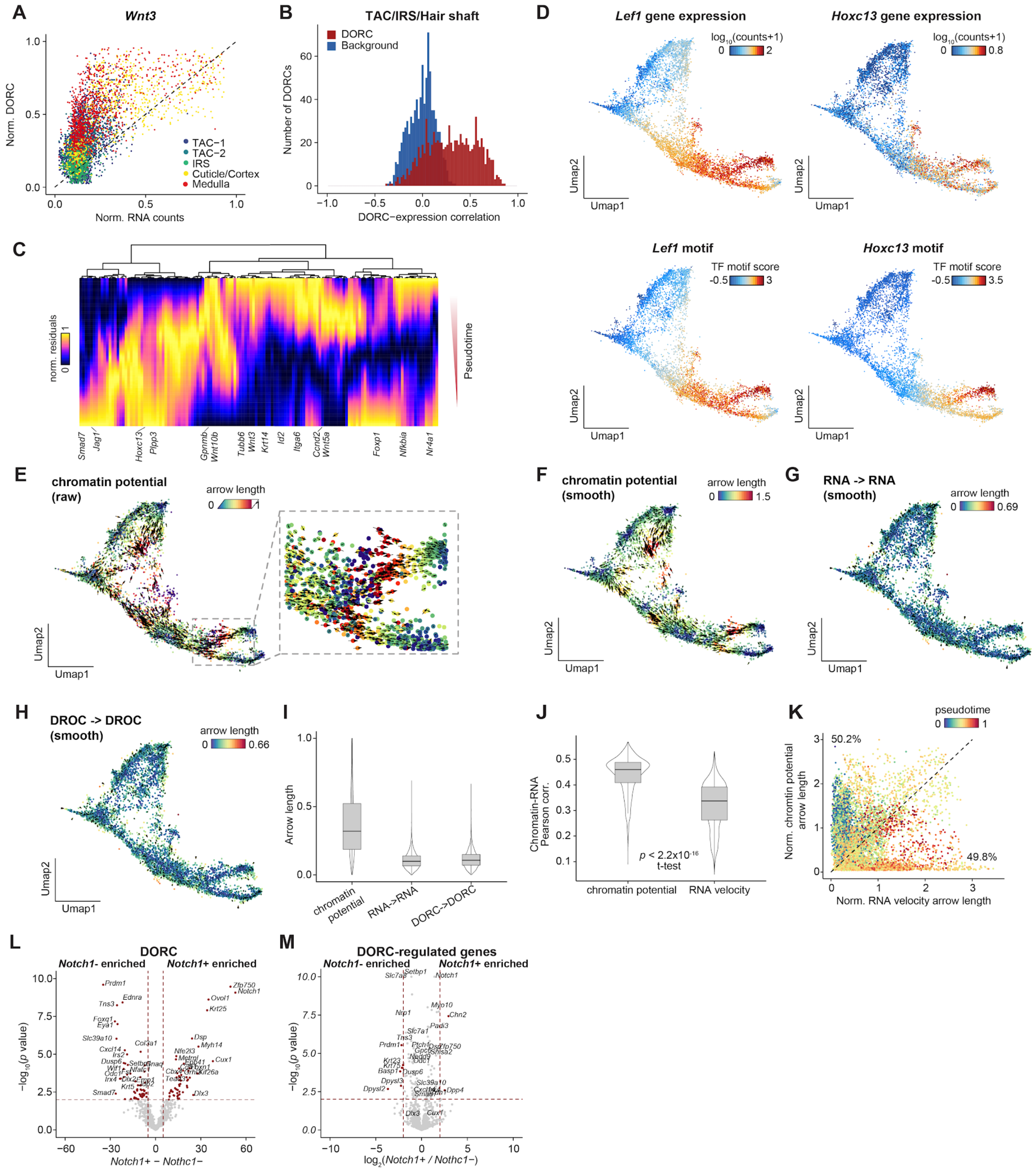

**Figure S8. Characterization of lineage-priming and chromatin potential.**
(A) The scatter plot of the *Wnt3* DORC score and *Wnt3* gene expression.
(B) The distribution of Spearman correlation between DORC and DORC-regulated genes. A background correlation was calculated by permuting peaks with matched GC-content and accessibility.
(C) Normalized residuals between chromatin accessibility and gene expression for hair-shaft lineages.
(D) ATAC UMAP visualization of gene expression (top) and motif score (bottom) inferred from ATAC-seq.
(E) The raw chromatin potential. The arrow denotes the distance between a cell in chromatin accessibility space to its most similar cell in RNA space.

(F) Raw chromatin potential was smoothed by averaging 15 k-nearest neighbors for each given cell.
(G) The arrows denote the potential "future" RNA state (observed in another cell) which is best predicted by the current RNA state. The arrows show the most correlated neighbour in RNA space for a given cell in RNA space.
(H) The arrows denote the potential "future" chromatin state (observed in another cell) which is best predicted by the current chromatin state. The arrows show the most correlated neighbour in chromatin space for a given cell in chromatin space.
(I) Comparison of the arrow lengths between chromatin potential, RNA-RNA prediction and chromatin-chromatin prediction.
(J) The Pearson correlations between chromatin state of a cell and the potential "future" RNA state of the given cell, predicted by either chromatin potential (left) or RNA velocity (right).
(K) A scatter plot shows the differences in arrow length between chromatin potential or RNA velocity. The dot color denotes pseudotime.
(L) Volcano plot of differentially enriched DORCs between *Notch1*+ and *Notch1*- lineage-prime cells.
(M) Volcano plot of differentially enriched DORC-regulated genes between *Notch1*+ and *Notch1*- lineage-priming cells.

**Supplementary tables**

Table S1. Oligo design.

Table S2. Marker genes for each cluster in mouse skin.

Table S3. List of peak-gene associations identified in GM12878 cells.

Table S4. List of peak-gene associations identified in mouse skin.

Table S5. DORCs overlap with known super-enhancers.
